# Reduction of neuronal activity mediated by blood-vessel regression in the brain

**DOI:** 10.1101/2020.09.15.262782

**Authors:** Xiaofei Gao, Jun-Liszt Li, Xingjun Chen, Bo Ci, Fei Chen, Nannan Lu, Bo Shen, Lijun Zheng, Jie-Min Jia, Yating Yi, Shiwen Zhang, Ying-Chao Shi, Kaibin Shi, Nicholas E Propson, Yubin Huang, Katherine Poinsatte, Zhaohuan Zhang, Yuanlei Yue, Dale B Bosco, Ying-mei Lu, Shi-bing Yang, Ralf H. Adams, Volkhard Lindner, Fen Huang, Long-Jun Wu, Hui Zheng, Feng Han, Simon Hippenmeyer, Ann M. Stowe, Bo Peng, Marta Margeta, Xiaoqun Wang, Qiang Liu, Jakob Körbelin, Martin Trepel, Hui Lu, Bo O. Zhou, Hu Zhao, Wenzhi Su, Robert M. Bachoo, Woo-ping Ge

## Abstract

The brain vasculature supplies neurons with glucose and oxygen, but little is known about how vascular plasticity contributes to brain function. Using longitudinal *in vivo* imaging, we reported that a substantial proportion of blood vessels in the adult brain sporadically occluded and regressed. Their regression proceeded through sequential stages of blood-flow occlusion, endothelial cell collapse, relocation or loss of pericytes, and retraction of glial endfeet. Regressing vessels were found to be widespread in mouse, monkey and human brains. Both brief occlusions of the middle cerebral artery and lipopolysaccharide-mediated inflammation induced an increase of vessel regression. Blockage of leukocyte adhesion to endothelial cells alleviated LPS-induced vessel regression. We further revealed that blood vessel regression caused a reduction of neuronal activity due to a dysfunction in mitochondrial metabolism and glutamate production. Our results elucidate the mechanism of vessel regression and its role in neuronal function in the adult brain.

## Introduction

Microvessels are composed of two closely interacting cell types, namely endothelial cells and pericytes (Armulik et al., 2011), that are embedded within the endfeet of astrocytes (Zhang et al., 1999). Neurons, astrocytic endfeet, pericytes, and endothelial cells as well as the vascular basal laminar layers form the neurovascular unit that coordinates and maintains brain metabolic activity and neuronal function (Abbott et al., 2006; Barres, 2008). Dysfunction or dysregulation of the neurovascular unit is associated with many diseases including stroke and Alzheimer’s disease (Iadecola, 2017; Zlokovic, 2011).

The density of brain vasculature and neurons declines significantly during normal aging (10–30%), and this decrease reaches 40–60% for Alzheimer’s patients (Bell and Ball, 1981; Brown and Thore, 2011; Riddle et al., 2003). However, little is known about how blood-vessel dynamics change with age and how this affects the activity of the adult mammalian brain(Chow et al., 2020; Coelho-Santos and Shih, 2020; Kaplan et al., 2020). Vessel regression was reported nearly two centuries ago (Brown and Thore, 2011) and has been studied in the vasculature of embryonic and postnatal rodents (Franco et al., 2015; Hughes and Chang-Ling, 2000; Korn et al., 2014; Rao et al., 2007) and zebrafish (Chen et al., 2012; Lenard et al., 2015). Several molecules (e.g., Wnt, Angiopoietin, Nrarp, VEGF, etc.) are found to be involved in normal vessel regression in developing organs (Baffert et al., 2006; Hammes et al., 2004; Inai et al., 2004; Lobov et al., 2005; Phng et al., 2009), However, the mechanisms underlying vessel regression are largely unknown, as are the fates of the various cellular components of the neurovascular unit. Here, using genetic methods to specifically label endothelial cells, pericytes, and glial cells with different fluorescent proteins, we performed longitudinal *in vivo* imaging of functional microcirculation for up to 6 months, enabling a comprehensive understanding of vessel regression as the brain develops and ages and how vessel regression–related changes in the microcirculation contribute to neuronal activity in the adult brain.

## Results

### Dysfunction of the microcirculation in the adult mouse brain

To evaluate the plasticity of the blood vasculature in the adult brain, we measured functional blood flow after administering FITC-dextran (500 kDa) to normal mice via the tail vein (**Figure 1A**). Blood flowrate was assessed by counting blood cells passing through the vessel. With this strategy, we monitored blood circulation within the same region of the cerebral cortex weekly for 3–6 months (**Figure 1B**). Surprisingly, we found that 1.72% of the microvessels became non-functional (i.e., no FITC signal) across the entire field within a 5-week window (**Figure 1C**). Interestingly, blood flow to ~75% of the dysfunctional microvessels, which were observed at day 1, was restored within a week (76.19%, n = 16 of 21). In the remaining vessels (i.e., occluded vessels), reperfusion was undetectable for >2 months after occlusion was first initiated, likely reflecting permanent functional loss (**Figure 1B, D, E**). These results demonstrated that a substantial proportionof blood vessels in the brain become dysfunctional under physiological conditions.

**Figure 1.**
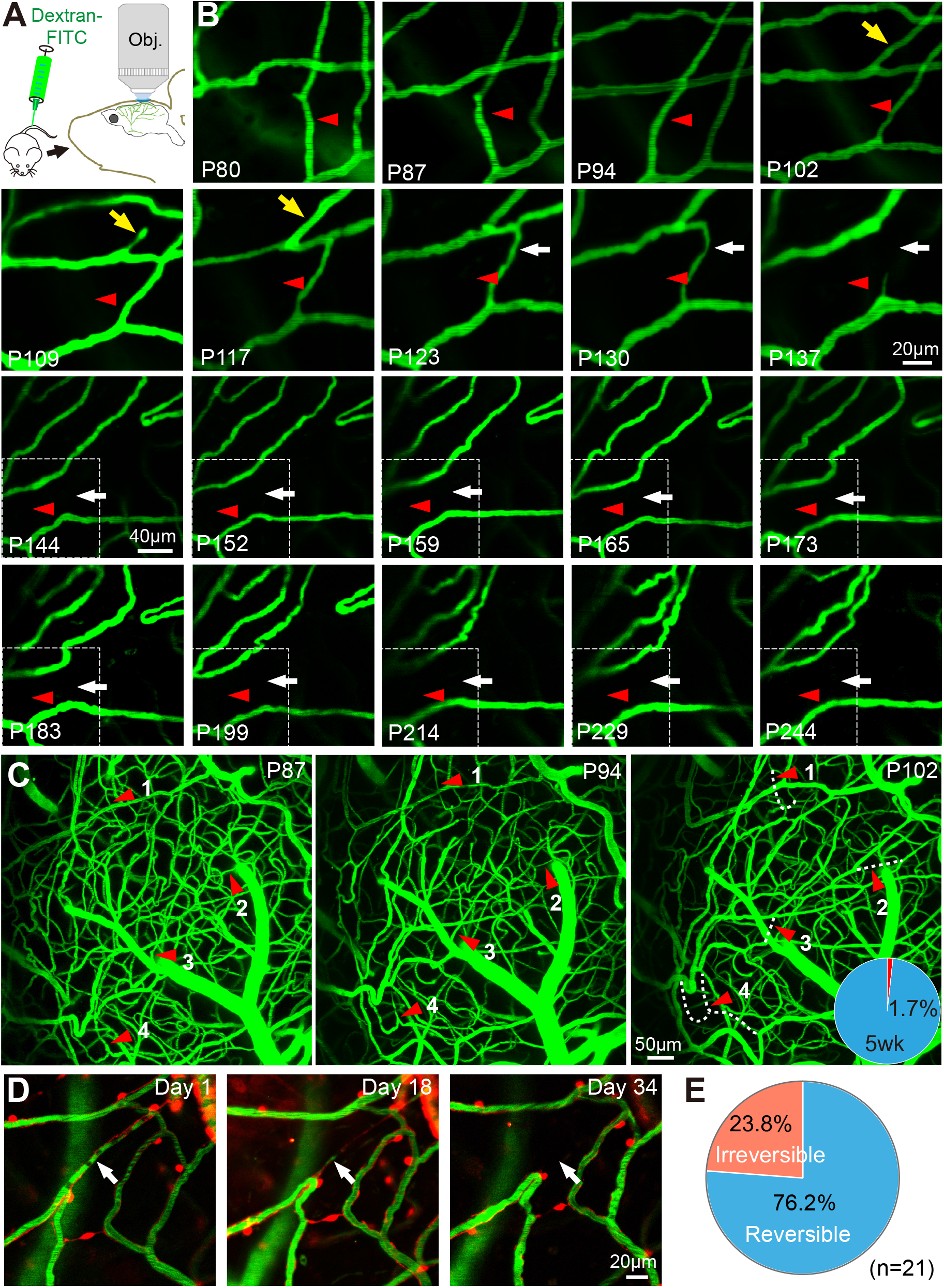
Dysfunction of the microcirculation in the adult mouse brain. (**A**) Schematic showing injection of dextran-FITC into the bloodstream via the tail vein and live imaging of functional brain microcirculation through a cranial window. (**B**) Time-lapse imaging of the microcirculation of a cortical region in a mouse from its postnatal day (P)80 to P244. The same region was imaged once every week for 6 months. Two different vessels (white arrows and red arrowheads) became dysfunctional, and blood flow was not restored after 102 and 137 days (i.e., which were 107 and 142 days after occlusion, respectively). Blood flow in one occluded blood vessel was restored (yellow arrows). From day144, each image has a larger field than prior images. (**C**) One example of images of all dysfunctional vessels in the field imaged. The lengths of these dysfunctional vessels (white dashed lines, 1 to 4) were used to normalize all vessels in the field. The percentages of dysfunctional blood vessels, which were normalized to all blood vessels in terms of length, are shown in the pie chart in the right panel. 5wks, time-lapse imaging of the region shown in (c) over 5 weeks. (**D**) Blood-flow occlusion precedes vessel regression. DsRed, pericytes in the brain of *NG2DsRedBAC* transgenic mice; green, FITC-dextran signal in blood vessels. Arrows indicate a regressing blood vessel. (**E**) Summary of results for blood vessels with or without blood-flow restoration (i.e., reperfusion) within a week after detecting occlusion. n = 21: number of occluded blood vessels imaged.

To determine whether loss of reperfusion in the microcirculation led to vessel regression, we used transgenic mouse strain *NG2DsRedBAC* (Zhu et al., 2008) for live imaging of pericytes via labeling with the fluorescent protein DsRed (**Figure 1D, Video 1**), which is specific for pericytes in veins, capillaries, and smooth-muscle cells in arteries/arterioles in adult mouse brains (**Figure S1**)(Hill et al., 2015; Jia et al., 2017). We detected that occlusion of blood flow for >1 week resulted in the disappearance of blood vessels (100%, n = 8 regressing vessels, **Figure 1D, Video 2–4**). Based on our staining results for different vascular cell types (**Figure 2–4**), loss of pericytes indicated near-complete vessel regression in all cases.

**Figure 2.**
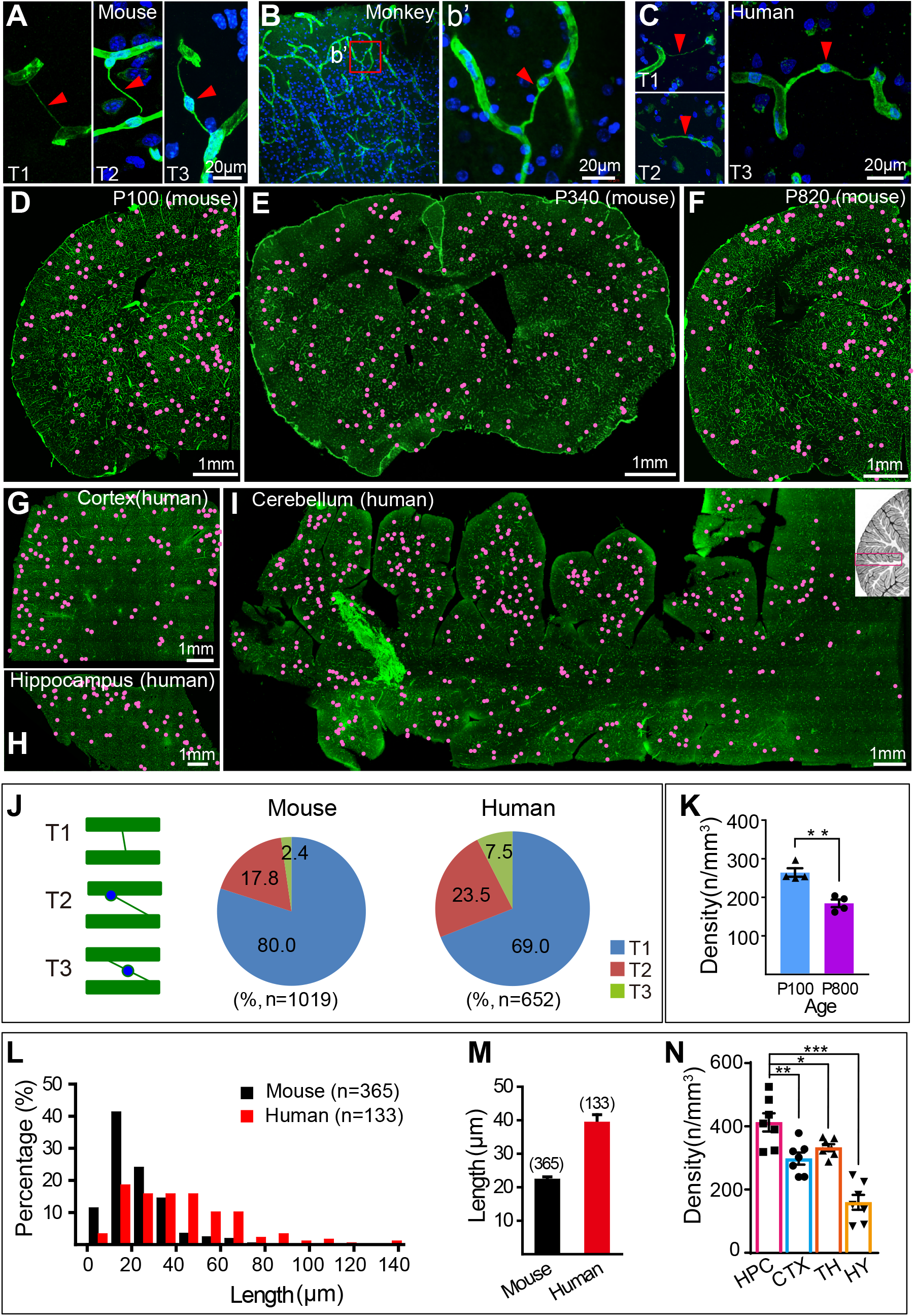
Distribution and properties of regressing vessels in the adult brain of different mammals. (**A**) Three typical types of regressive vessel structures in the adult mouse brain (T1, T2, T3). Blue, nuclei stained with Hoechst 33342; green, anti-laminin; arrowheads, regressing vessels. (**B**) Regressing vessels detected in the cortical section of a 3-year old monkey. (**C**) Three types of representative regressing vessels from human brain tissues. (**D–F**) Distribution of all regressing vessels in whole-brain/hemisphere sections of mice at P100, P340, and P820. Each dot represents one regressive vessel. All vessels were labeled with anti-laminin or anti-collagen IV. (**G–I**) Distribution of regressive vessels detected in three different brain regions of a 45-year-old human male: cerebral cortex (G), hippocampus (H), cerebellum (I). The location of the imaged section from the cerebellum is shown in the top right corner in (i) outlined in red. All vessels were labeled with anti-laminin. Slice thickness, 70 *μ*m. (**J**) Schematic diagrams and percentages of three different types of regressive structures found in human and mouse brains. Green, blood vessels; blue, soma. (**K**) Density of regressive vessels in the brain of a young adult (~P100) and old mouse (~P800). (**L**) Length distribution of regressing vessels from human and mouse brains. The *x* axis represents the length of regressing vessels. (**M**) Average length of regressing vessels in human and mouse brains. (**N**) Comparison of the density of regressing vessels from different mouse brain regions. HPC, hippocampus; CTX, cerebral cortex; TH, thalamus; HY, hypothalamus. \**p* < 0.05; \*\**p* < 0.01, \*\*\**p* < 0.001, two-tailed unpaired *t*-test. All error bars indicate SEM.

### Distribution and properties of regressing vessels in the adult brain of different mammals

Endothelial cells and pericytes are completely enwrapped by a sheath comprising the extracellular matrix molecules laminin and collagen IV(Armulik et al., 2011). Antibodies specific for collagen IV and laminin have been used to stain regressive vessels in the brains of different species(Brown, 2010; Cammermeyer, 1960), but the distribution of regressing vessels throughout the mammalian brain has never been determined. Labeling blood vessels with anti-laminin or anti-collagen IV revealed three distinct regressive vessel types in mouse, monkey, and human brains (**Figure 2**). The first type (Type I, T1) constituted 70–80% of all regressive vessels and appeared as a thin, string-like process (1–2 μm) connecting two neighboring blood vessels; no soma was associated with this type (**Figure 2A, C**). The second type (Type II, T2) included a single soma situated at one end of the process, adjacent to neighboring blood vessels (**Figure 2A, C**). The third type (Type III, T3) had a single soma located in the middle of the laminin-positive process with both ends of its processes connecting neighboring blood vessels (**Figure 2A, C**). The abundance of these three types of regressing vessels was similar in human and mouse brain sections (**Figure 2J**) (Types I, II, III in humans were 69.0%, 23.5%, and 7.5%, respectively, n = 652; in mouse, 80.0%, 17.8%, and 2.4%, n = 1019; **Figure 2J**).

To understand how the distribution of regressive vessels in the rodent brain varies with age, mouse whole-brain sections were scanned at high resolution (**Figure S2**), and tiled images were analyzed for each brain section (**Figure 2D–F, Video 5**). The same strategy was used to detect regressing vessels in several human brain regions including the hippocampus, cerebellum, and cerebral cortex (**Figure 2G–I**). The regressing vessels were widely distributed in nearly all major brain regions in mice (**Figure 2D–F, Video 5**), and the length of regressive vessels did not differ between juvenile and adult mouse brains (P17, 22.95 ± 0.82 μm, n = 227; P340, 21.72 ± 1.18 μm, n = 138). Interestingly, there was a significant decrease in the density of regressing vessels in the aging brains, but the regressive vessels were still abundant in aged brains (P100, 262.3 ± 11.1/mm^3^, n = 4 mice; P800–820, 177.9 ± 12.6/mm^3^, n = 4 mice; **Figure 2K**). On average, the regressing vessels were significantly longer in the adult human brain than in mouse brain (human, 39.4 ± 2.2 μm, n = 133; mouse, 22.4 ± 0.9 μm, n = 365, **Figure 2L, M**). Interestingly, the hippocampus had the highest density of regressing vessels among all the brain regions we assessed (**Figure 2N**).

### Mechanism of blood-vessel regression in the brain

Although blood-vessel regression has been studied for decades (Brown, 2010), the underlying cellular mechanism remains controversial, including the fate of pericytes. Pericytes enwrap endothelial cells(Armulik et al., 2011), but it is unclear whether regressive vessels form with laminin alone, also known as ghost structures or string vessels labeled by anti-collagen IV or anti-laminin (Brown, 2010; Challa et al., 2002), or whether they include endothelial cells and/or pericytes. An analysis of the staining (anti-CD31 and anti-laminin/anti-collagen IV) of the cellular constituents of ~200 regressive vessels in brain sections from *NG2DsRedBAC* mice revealed the presence of pericytes in ~90% of remnants of regressive vessels (**Figure 3A–C**). Similar results were obtained with brain sections of *Pdgfrb-Cre::Ai14* (**Figure S3**), which is a specific mouse strain for labeling pericytes in the brain (Cuttler et al., 2011). Capillaries are composed of laminin layers, endothelial cells, and pericytes(Armulik et al., 2011), implying seven distinct possible combinations of cell components for any given regressive vessel. Only three of these combinations were found in the regressive structures that we analyzed (**Figure 3C**): a very small percentage of regressive vessels were laminin^+^ DsRed^+^CD31^+^ (5.6%, 10 of 186, i.e., pericytes, laminin layer and endothelial cells) or laminin^+^ only (9.2%, 17 of 186, i.e., laminin layer only), and the remainder contained laminin layers and pericytes but no endothelial cells (laminin^+^DsRed^+^CD31^-^, 85.2%, 159 of 186). These results indicated that endothelial cells are eliminated prior to the loss of pericytes during vessel regression. This differs from previous reports that concluded that string-like vessels (i.e. regressing vessel) are “ghost” vessels comprised only of laminin matrix(Brown, 2010). Our results indicate that >90% of regressing vessel structures contained laminin- or collagen-positive layers and pericytes (**Figure 3C**). These findings were supported by anti-collagen IV staining of brain sections of *Cdh5-CreER::Ai14* mice (**Figure S4, S5**), which is a specific mouse strain for labeling endothelial cells with tdTomato in the brain. We detected very few regressive vessels that contained endothelial cell component (7.59%, n = 28 of 369 regressive vessels). This was verified by *in vivo* imaging results from *Cdh5-CreER::Ai6::NG2DsRedBAC* mice, in which the endothelial cells (Sorensen et al., 2009) and pericytes were labeled with ZsGreen and DsRed, respectively (**Figure S5**). These results indicate the endothelial cells initially retracted in regressing vessels, after which pericytes were retained beyond the 3-week time point (**Figure S5**).

**Figure 3.**
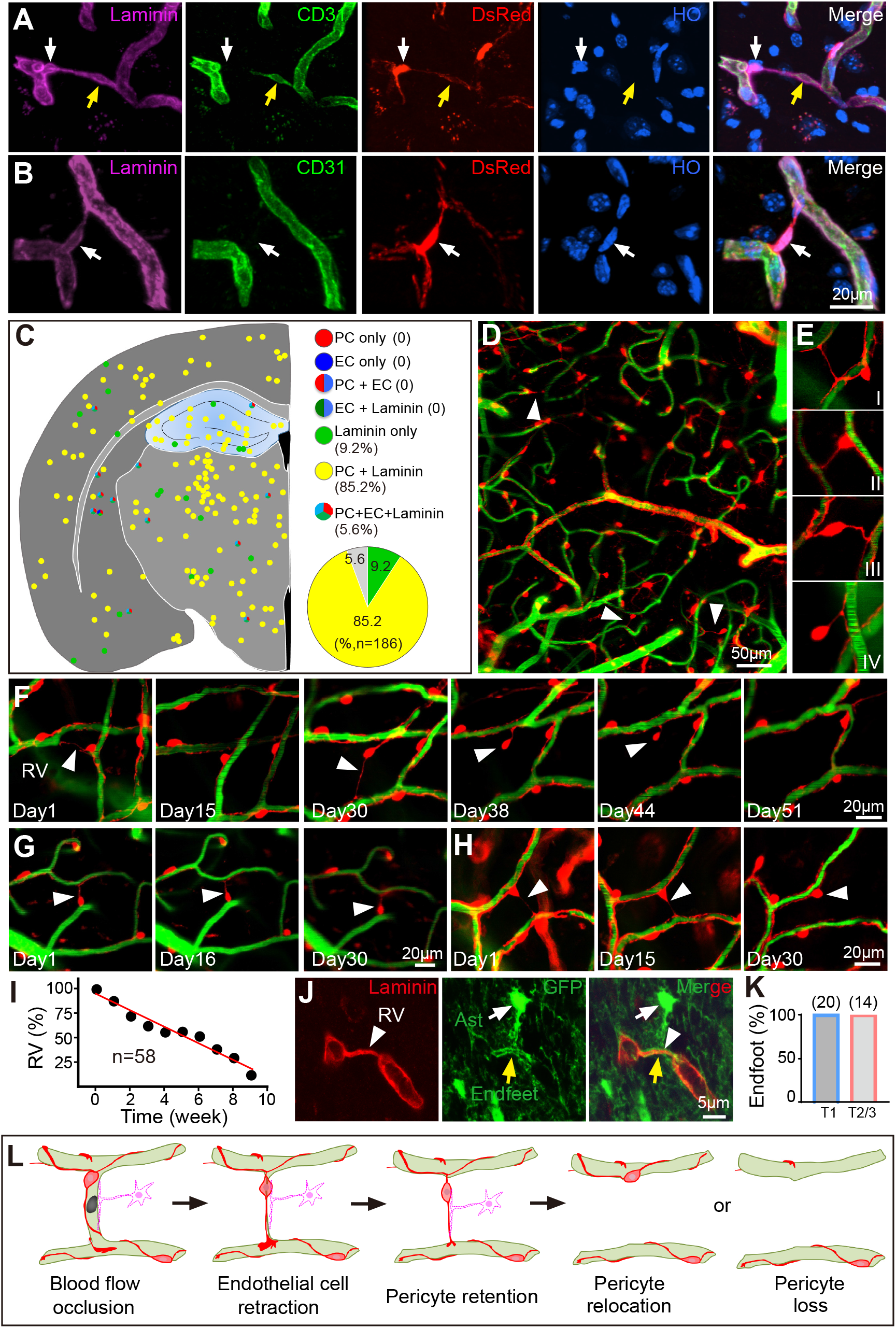
Mechanism of blood-vessel regression in the brain. (**A**, **B**) Cellular components of regressing vessels in the brain. Magenta, laminin layer stained with antilaminin; green, endothelial cells labeled with anti-CD31; red, DsRed signal for labeling pericytes in a *NG2DsRedBAC* transgenic mouse; blue, nuclei stained with Hoechst 33342 (HO). (**C**) Percentages of regressing blood vessels with different cell components: PC, pericytes; EC, endothelial cells; laminin, laminin layer. Inset, seven possible combinations of cell components. (**D**, **E**) *In vivo* imaging of regressing vessels in the brain of an adult *NG2DsRedBAC* mouse. Four types of regressing vessels were observed *in vivo*: I, the soma of the pericyte was located in a neighboring vessel; II, the soma of the pericyte was located at one end of the regressive vessel; III, the soma was located in the middle of the regressive vessels; IV, one end of the regressive vessel had detached from the neighboring blood vessel. (**F–H**) Time-lapse *in vivo* imaging of the entire process of vessel regression and pericyte fate in regressing vessels. Three different fates of pericytes (arrowheads): cell death (**F**), retention at the same location (**G**), and relocation to a neighboring vessel (**H**). Blood flow was assessed by labeling with dextran-FITC-500K. Pericytes (red, DsRed) were labeled in the transgenic mouse line *NG2DsRedBAC*. (**D–H**). The examples in (**F**) and Figure 1 (**B**) were from the same field of a mouse. (**I**) Summary of lifespan for 58 regressing blood vessels that were imaged. (**J**) Astrocytic endfeet around regressing vessels. Green, GFP expressed in astrocytes from *hGFAP-GFP* transgenic mice. Endfeet (yellow arrows) of astrocytes (Ast, white arrows) enwrapped the entire surface of regressing vessels (RV, white arrowheads), which were stained with anti-laminin (red). (**K**) Percentages of regressing vessels fully enwrapped by astrocytic endfeet. T1–3 indicate three distinct types of regressing vessels. (**L**) Our model of vessel regression in the adult brain. Vessel regression starts with blood-flow occlusion (dark region inside the vessel) in a certain percentage of blood vessels. Endothelial cells (light green) retract rapidly in response to occlusion. Pericytes (red) remain for a long period and form a typical regressive structure with a laminin layer. Pericytes in regressing blood vessels either relocate to neighboring blood vessels or die.

To investigate pericyte fate during vessel regression in the adult brain. We imaged pericytes in regressing vessels from cortical regions of *NG2DsRedBAC* mice. Blood flow was monitored with FITC-dextran or with blood cells labeled with 3,3’-dioctadecyloxacarbocyanine perchlorate (commonly called DiO). The morphology of pericytes in regressing vessels varied, as assessed *in vivo* (**Figure 3D, E**), which represented different stages of vessel regression observed in fixed tissues (**Figure 2A–C**). Longitudinal live imaging revealed that the half-life of regressing vessels was ~5 weeks (n = 58, **Figure 3F–I**). Pericytes from regressing vessels had three distinguishable fates: ~30% disappeared, likely by apoptosis (**Figure 3F**); 20% remained stable, with no obvious change in morphology or somatic location over the 8-week period (**Figure 3G**); the remaining ~50% retracted one of their processes, and the soma relocated to a neighboring blood vessel (**Figure 3H**). These results indicated that, in comparison with endothelial cells, pericytes in regressing vessels maintained a relatively stable structure. To further verify these results, we took advantage of a mosaic analysis with double markers (*MADM*) transgenic mouse line (Zong et al., 2005) for labeling single pericytes through breeding them with *Hprt-Cre* mice in which the Cre cassette is inserted into the X-linked gene *Hprt*. These mice have Cre recombinase in their oocytes, which excises a floxed sequence at the zygote or early cleavage stage (Tang et al., 2002). In *Hprt-Cre::MADM-7GT* mice (Hippenmeyer et al., 2013), brain cells, including pericytes, were sparsely labeled with GFP and/or DsRed (**Figure S6A**), allowing high-resolution imaging of individual pericytes of regressing vessels (**Figure S6**). Surprisingly, these single pericytes in regressing blood vessels had two mature and complex processes that wrapped around adjacent blood vessels (n = 8), unlike regular/typical pericytes (**Figure S6**), both complex processes are connected with a string structure from the same pericyte, which is consistent with our results from long-term *in vivo* imaging in *NG2BACDsRed* mice (**Figure 3**). These results indicated that remnant pericytes of regressing vessels were relatively stable constituents of regressing vessels, which is consistent with our *in vivo* time-lapse imaging of vessel regression (**Figure 3F**).

Astrocytes are the most prevalent glial cells in the mammalian brain. They extend two types of processes from their cell bodies: fine perisynaptic processes that cover most neuronal synapses, and larger processes, known as endfeet, that primarily extend to and make tight contacts with blood vessels, thus covering >99% of the abluminal vascular surface in the adult brain(Zhang et al., 1999). Astrocytic endfeet ferry glucose from blood to neurons and are vital for neuronal function(Barres, 2008; Iadecola and Nedergaard, 2007). Despite being one of the critical cells of the neurovascular unit, the fate of astrocytic endfeet in vessel regression is unclear. Therefore, we stained brain sections of *hGFAP-GFP(Zhuo et al., 1997*) or *Aldh1L1-EGFP(Gong et al., 2003*) transgenic mice with anti-laminin or anticollagen IV to locate regressing vessels, all of which were fully enwrapped by astrocytic endfeet (100%, n = 34 of 34, **Figure 3J, K**). These results indicated that vessel regression was not initiated by the retraction of astrocytic endfeet. Overall, our results demonstrate that vessel regression in mammals is mediated by distinct steps: blood flow is occluded, endothelial cells are reabsorbed, and pericytes relocate or die; finally, glial endfeet lose contact with vessels (**Figure 3L**).

### Short-term arterial occlusion induces an increase in vessel regression in the adult brain

Since all vessel regressions that we detected occurred after vessel occlusion (Figure 1), we wondered whether manipulating blood flow by introducing a brief middle cerebral artery (MCA) occlusion would lead to an increase of vessel regression. It has been shown that MCA occlusion lasting less than 20 min does not cause neuronal cell death (Carmichael, 2005; Kokaia et al., 1995), but can induce an inflammatory response in the brain (del Zoppo, 2010). Furthermore, upregulation of inflammation under chronic cerebral hypoperfusion and Alzheimer’s disesase induced stagnation of leukocytes in blood vessels (Cruz Hernandez et al., 2019; Yata et al., 2014). We therefore performed short-term (15 or 20 min) occlusion of the MCA to induce inflammation in adult mouse brains and subsequently assessed the density of vessel regression (**Figure 4A**). We observed that a 20-min MCA occlusion significantly increased the density of regressive blood vessels in the occluded hemisphere 15 d after occlusion (156.90 ± 12.07%, n = 5 mice, **Figure 4B–D**). We also observed a slight increase of blood vessel density in the occluded hemisphere (103.02± 4.83%, n = 5 mice, **Figure 4D**), which was due to angiogenesis after MCA occlusion. To determine whether the observed regression was due to pruning of newly formed vessels, we measured the density of regressing vessels 7 d after 15-min occlusion. We observed that angiogenesis had not occurred in the occluded hemisphere at this time point (100.53 ± 2.44%, n = 6 mice, **Figure 4E–G**). Under this condition, we detected that the density of regressing vessels increased by ~50% in the hemisphere with MCA occlusion (151.08 ± 9.72%, n = 6 mice, **Figure 4E, F, H**). The results demonstrate that a brief occlusion induces a rapid, dramatic increase of vessel regression in the adult brain and that these regressing vessels derive from the existing blood vessels rather than newly formed ones.

**Figure 4.**
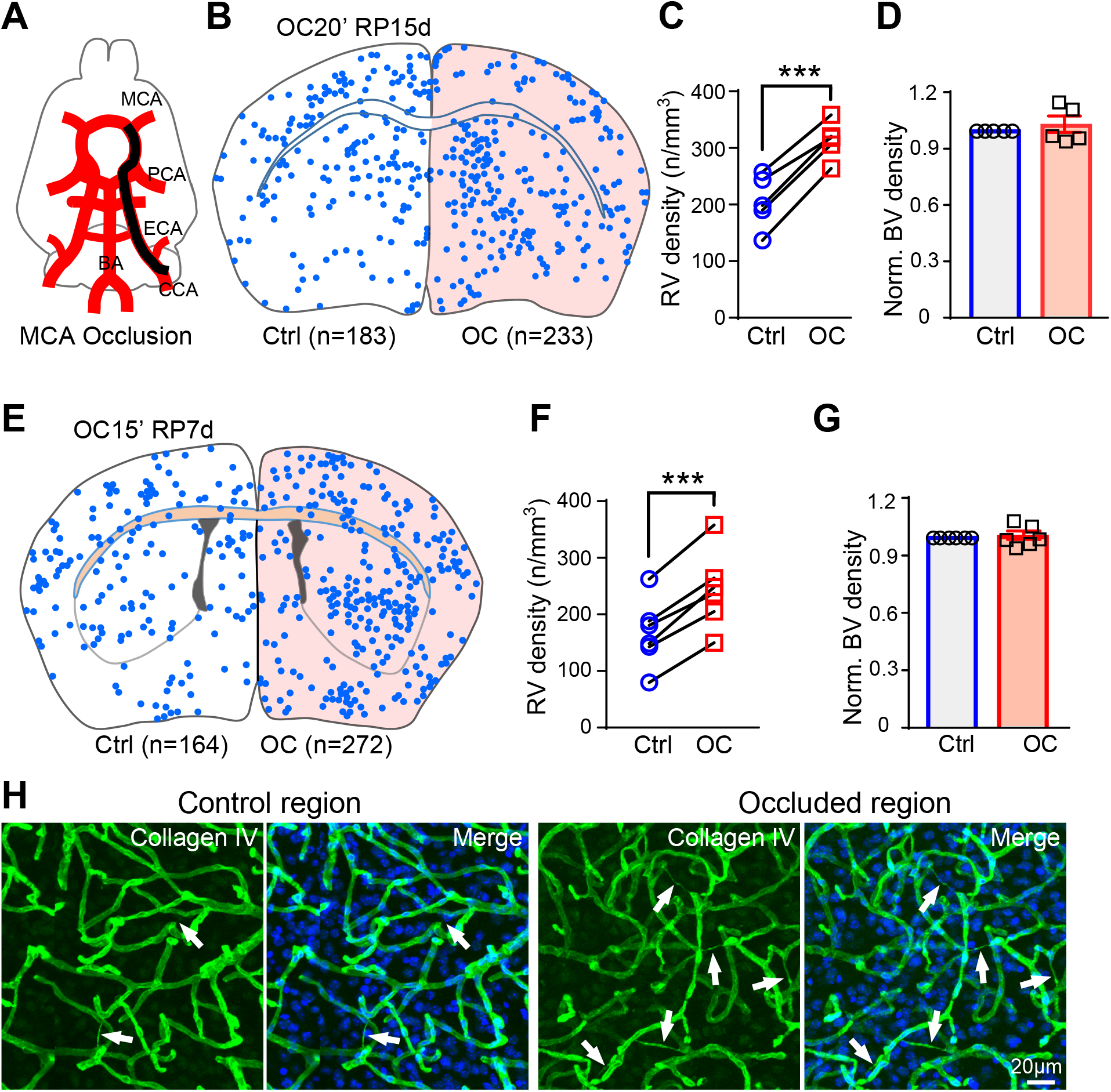
Short-term arterial occlusion induces an increase in vessel regression in the adult brain. (**A**) Schematic illustrating the brief middle cerebral artery (MCA) occlusion was briefly occluded (15 or 20 min). (**B**) Distribution of regressing vessels in a whole-brain section stained with anti-collagen IV after MCA occlusion. Each blue dot represents one regressive vessel. OC20’RP15d indicates 20 min occlusion with subsequent 15 d reperfusion. OC, hemisphere with MCA occlusion; Ctrl, hemisphere with no MCA occlusion. (**C**) Quantitation of the density of regressive vessels in the left (Ctrl) and right (OC) hemispheres. Dots on the left and right sides represent the density of regressing vessels from Ctrl (blue) and OC hemispheres (red), respectively. ***, *p* < 0.001. Paird *t*-test. All error bars indicate s.e.m. (**D**) Normalized blood vessel densities in Ctrl and OC hemispheres. (**E**) Distribution of regressing vessels in a whole-brain section after MCA occlusion. The mouse was perfused and blood vessels were stained with anti-collagen IV 7d after MCA occlusion. OC15’RP7d, 15 min occlusion with subsequent 7 d reperfusion. (**F**) Quantitation of the density of regressive vessels in the left (Ctrl) and right (OC) hemispheres. Dots on the left and right sides represent the density of regressing vessels from Ctrl (blue) and OC hemisphere (red), respectively. ***, *p* < 0.001. Paird *t*-test. All error bars indicate s.e.m. (**G**) Normalized blood vessel densities in Ctrl and OC hemispheres. (**H**) Representative images demonstrating blood vessel pattern and regressive vessels (arrows) in Ctrl and OC hemispheres. Green, anti-collagen IV; blue, nuclei stained with Hochest33342.

### Inflammation induces a dramatic increase in vessel regression in the adult brain

To further investigate whether inflammation outside of the brain also causes vessel regression in the brain, and whether leukocyte adhesion in blood vessels leads to vessel occlusion and subsequently to vessel regression, we intraperitoneally injected lipopolysaccharide (Huang et al.), a component of the cell walls of Gram-negative bacteria, which is commonly used in the laboratory to induce a pro-inflammatory response. After we injected mice with a low dose of LPS daily for four days, a higher density of leukocytes could be detected in the cerebral blood circulation. Twenty-one days after the first injection of LPS, we stained brain sections with anti-collagen IV and found the density of regressive blood vessels had increased dramatically compared to that of the control group that received PBS injections. Administration of Ly6G antibodies can impair surface expression of β2-integrins in inflammation-stimulated neutrophils and inhibit ICAM-1 binding (Wang et al., 2012). As a result, these antibodies can be used to prevent firm adhesion of leukocytes to activated endothelial cells. After we injected LPS, we administered Ly6G antibodies once every 3 days until the mice were euthanized on day 21. We observed that the density of regressive blood vessels in the Ly6G group was significantly reduced when compared to the LPS group (Figure 5C), indicating that LPS-mediated acute inflammation induces vessel regression in the brain and that this regression is mainly due to leukocyte stagnation.

**Figure 5.**
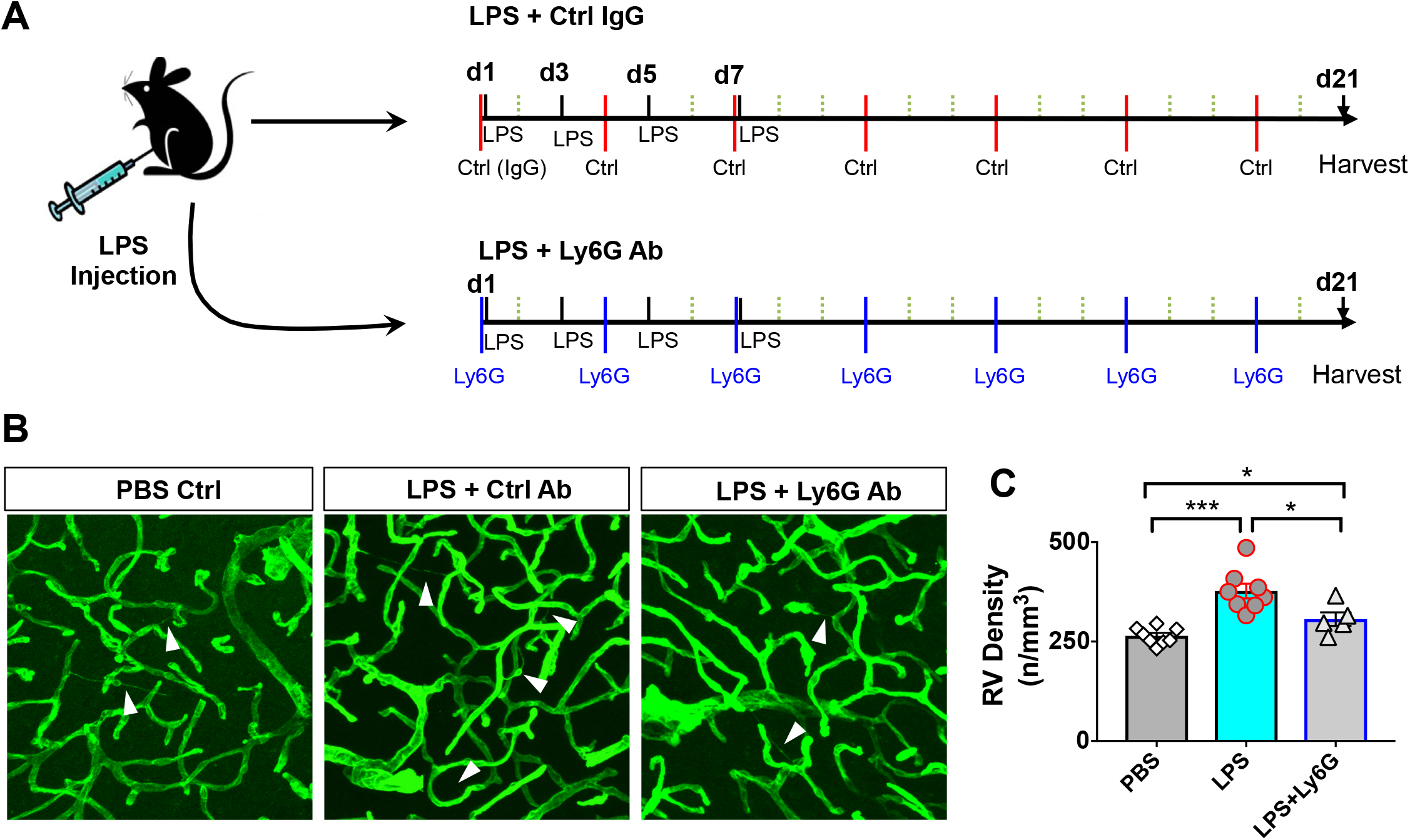
LPS-mediated inflammation induces an increase of brain vessel regression. (**A**). Schematic illustrating the administration of LPS and control or Ly6G antibodies before euthanizing mice for immunostaining. Antibodies were administered i.p. to 12- to 14-week-old mice on days 1, 4, 7, 10, 13, 16, and 19 of the study. LPS was injected on days 1, 3, 5, and 7. Mouse brains were collected for Collagen IV immunostaining on day 21. (**B**) Representative immunostaining results of blood vessels from the cortical regions of control and LPS-injected mice before and after administration of Ly6G antibodies. Arrowheads, regressive blood vessels. Green, staining for Collagen IV. (**C**) A summary of regressive blood vessel density (the number of regressive vessels normalized to the volume of brain tissue) in the brains of mice after injection of PBS or control antibodies (n = 8 mice), LPS and Ctrl IgG (n = 8 mice), or the combination of LPS and Ly6G antibodies (n = 5 mice). \**p* < 0.05; \*\*\**p* < 0.001, two-tailed unpaired Student’s *t*-test. All error bars indicate s.e.m.

### Reduction of neuronal activity was mediated by blood-vessel regression

In the mammalian brain, blood-vessel density decreases substantially with age (Brown, 2010; Riddle et al., 2003). Interestingly, we hardly detected newly formed functional blood vessels in the adult brain even when we imaged the same regions for months (**Figure 1**). Thus, continued vessel regression increases the distance between neurons and blood vessels. Because few vessels actually undergo regression in a short time window, (i.e., ~0.35% per week), it was difficult to predict which vessels might regress for the purpose of imaging neuronal activity in the same region for months. To assess how vessel regression affects brain function, we created a conditional knockout of *Tak1* in endothelial cells (Ridder et al., 2015), as Tak1 is reported to be involved in vessel regression (Ridder et al., 2015). *Tak1* knockout increased vessel regression by 6–8 fold at 1–3 weeks after administration of tamoxifen to *Cdh5-CreER:Tak1^fl/fl^* mice (*Tak1 CKO*, **Figure 6A, B**). This allowed us to possibly evaluate the effect of vessel regression on neuronal activity in a relative shorter time frame. To detect neuronal activity, we infected neurons with adeno-associated virus (AAV) expressing GCaMP (AAV-Camk2-GCamp6m) in cortical layers 2–4 (**Figure. 6C–F**) at one month before tamoxifen injection. Neuronal activity was measured before and after conditional knockout of *Tak1* in endothelial cells in conscious mice (**Figure 6C–K**). To ensure detection of subtle differences in activity before and after vessel regression, we imaged and compared the same neurons at different time points. We observed that neuronal activity declined significantly 1–2 weeks after the conditional knockout of *Tak1* (2.83 ± 0.26 spikes/min before tamoxifen administration vs. 1.43 ± 0.19 spikes/min after administration (n = 65 neurons; **Figure 6E, G, I, J**). The frequency of all spikes decreased in the *Tak1*-knockout group, indicating that vessel regression reduced neuronal activity. In control mice, however, neuronal activity was not affected by tamoxifen (or by the carrier solution, as a control; 2.60 ± 0.24 spikes/min vs. 2.76 ± 0.23 spikes/min, respectively; n = 45 neurons; **Figure 6F, H, K**, and **L**). These results indicated that vessel regression in the microcirculation mediated the reduction of neuronal activity with age in the cerebral cortex.

**Figure 6.**
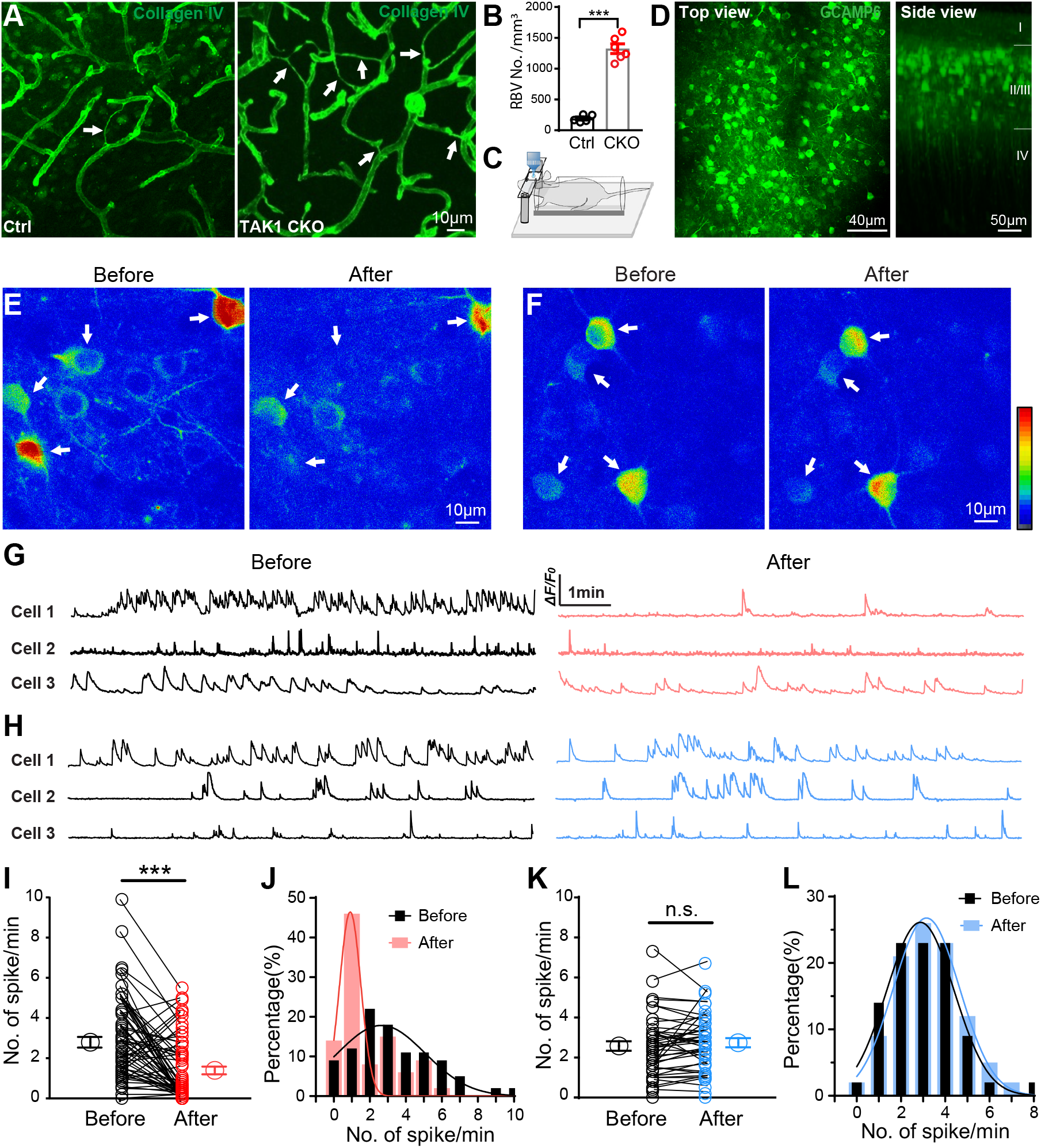
*In vivo* imaging of neuronal activity in *Cdh5-CreER:Tak1^fl/fl^* mice. (**A**) Example images of blood vessels in brain sections from control mice (Ctrl) or from *Cdh5-CreER:Tak1^fl/fl^* mice with a conditional knockout (Tak1 CKO) of *Tak1* in endothelial cells. (**B**) Statistical analysis of the density of regressive blood vessels in the brain of Tak1 CKO and control mice. \*\*\**p* < 0.001, Student’s *t*-test. (**C**) Strategy used to image neuronal activity in the brain of conscious mice. (**D**) GCaMP6 signal of neurons in layers II–IV of the cerebral cortex. AAV-Camk2-GCaMP6m was injected ~1 month before live imaging. I–IV indicates the layers of the cerebral cortex. (**E, F**) Example images of the GCaMP6 signal obtained from *Cdh5-CreER:Tak1^fl/fl^ (Tak1 CKO*) mice (E) and control mice (F) before and after administration of tamoxifen. Warm pseudocolor indicates a high calcium signal. (**G**, **H**) Representative traces of calcium transients recorded from three cortical neurons (arrows) of a *Tak1 CKO* mouse (G) and a control mouse (H) before and after a 1-week administration of tamoxifen. (**I**) Statistical analysis reveals the significance of differences of calcium transients in CKO mice as shown in (E) and (G). Each connected pair of black and red circles denotes data obtained with the same neuron before (black) and after (red) tamoxifen injection. Each of the two large circles denotes the mean value ± s.e.m. for the two groups. \*\*\**p* < 0.001, paired Student’s *t*-test. (**J**) Frequency distribution of calcium transients from all neurons before (black) and after (red) tamoxifen was administered. Red Gaussian curve shows that the spike frequency was shifted to the left (i.e., lower value) after injection of tamoxifen into *Tak1 CKO* mice. (**K**) Statistical analysis reveals no significant differences (n.s.) of calcium transients in *Tak1 CKO* mice as shown in (F) and (H). Each connected pair of black and blue circles denotes data obtained with the same neuron before (black) and after (blue) tamoxifen injection. Each of the two large circles denotes the mean value ± SEM for the two groups. (**L**) Frequency distribution of calcium transients from all neurons before (black) and after (blue) Tamoxifen solution was administered. The Gaussian curve shows that the spike frequency was not shifted after injection of tamoxifen into control mice (*Tak1^fl/fl^*).

### Vessel regression leads to abnormalities in neuronal metabolism and glutamate production

To determine whether the reduction of neuronal activity in the brains of *Tak1 CKO* mice was due to neuronal degeneration, we stained neurons with Fluro Jade C (Jia et al., 2017), a marker for all degenerating neurons regardless of specific insult or mechanism of cell death. We did not detect any degenerating neurons in brain sections of *Tak1 CKO* mice. Similar results were observed when we stained neurons with the antibodies against activated caspase-3, a marker for apoptotic cells. We further asked whether the accumulation of regressive vessels in *Tak1 CKO* results in mitochondrial dysfunction and affects energy generation in neurons. We performed electron microscopy (EM) to image brain sections from the cerebral cortex of control and *Tak1 CKO* mice. Mitochondria from neuronal synaptic terminals of the control and *Tak1 CKO* brains were compared, and we observed that mitochondria in the neurons of *Tak1 CKO* mice had abnormal morphology. Specifically, the cristae showed irregular morphologies and lower electrical signals under EM. The number of cristae decreased and they were unevenly distributed in the synaptic mitochondria of *Tak1 CKO* brains (**Figure 7A, B**).

**Figure 7.**
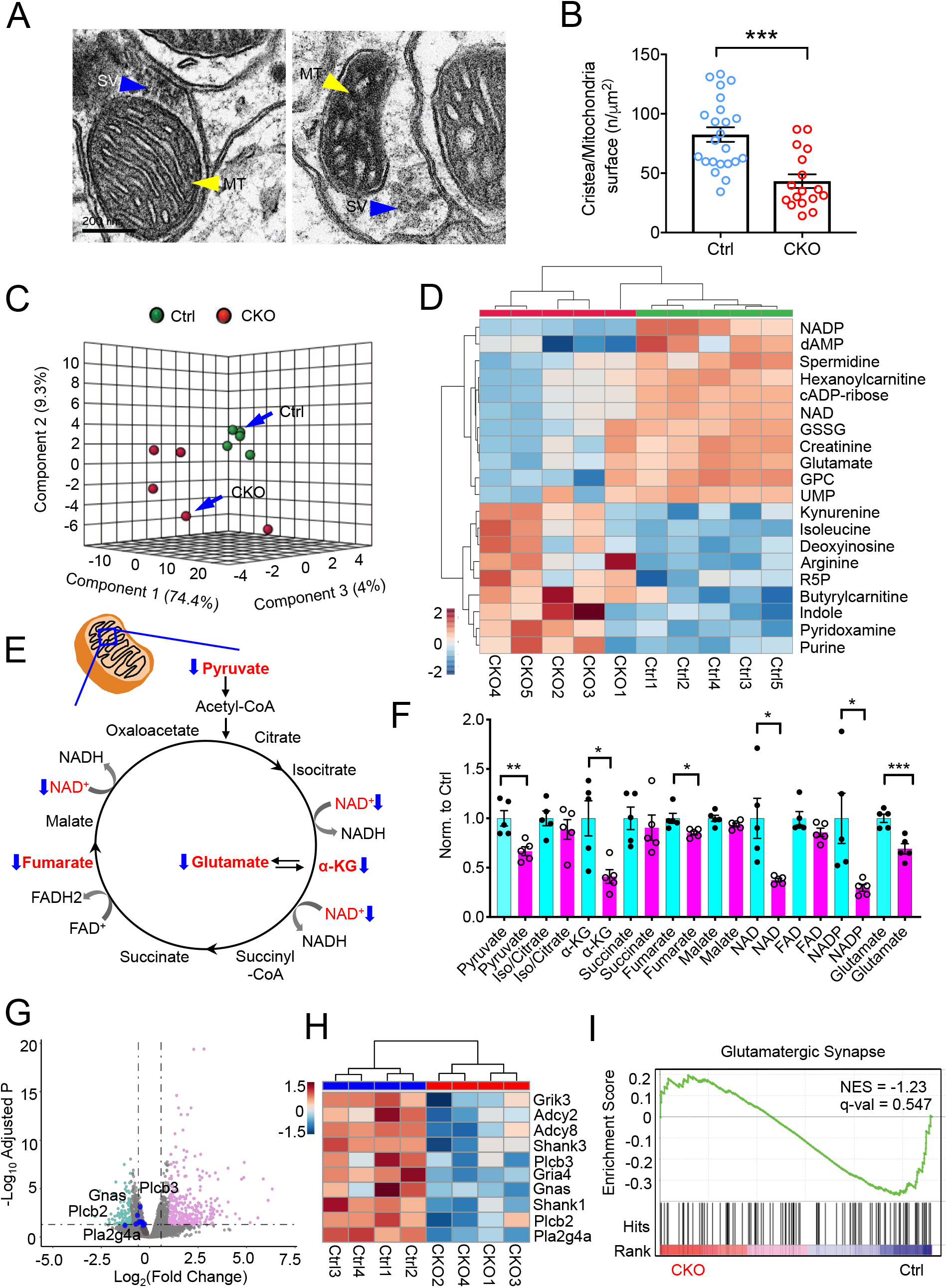
Abnormalities of neuronal metabolism in the brains of *Tak1 CKO* mice. (**A**) The morphology of mitochondria in neuronal synaptic terminals from control and *Tak1 CKO* mice (*Cdh5-CreER:Tak1^fl/fl^*). SV, synaptic vesicles. MT, mitochondria. (**B**) Summarized results of mitochondrial features in the neurons of control and *Tak1 CKO* mice. Y-axis, the number of cristae was normalized to the surface of mitochondria. ***, *p*<0.005, unpaired Student’s t-test. All error bars indicate s.e.m. (**C**). Principal component analysis (PCA) of the metabolomes of control (Ctrl) and *Tak1 CKO* brain samples. A total of 204 metabolites were measured from each sample and all detected metabolites were used for PCA. Three PCs explained 87.7% (74.4%, 9.3% and 4%) of the variance between Ctrl and *Tak1 CKO*. PC scores are indicated as %; circles indicate individual samples from the control and CKO groups. (**D**) A heatmap representation of 20 metabolites (VIP score > 1, i.e., Variable Importance in Projection, by partial least squares discriminant analysis) in the cerebral cortex of five control (Ctrl) and five *Tak1 CKO* mouse brains. All metabolites are listed on the right side of the map. Color bar (bottom left) indicates the scale of standardized metabolite levels. Warm color indicates higher concentration. NAD, Nicotinamide adenine dinucleotide; dAMP, deoxyadenosine monophosphate; GSSG, oxidized glutathione; GPC, Glycerophosphocholine; R5P, Ribose 5-phosphate. (**E**, **F**) Schematic (E) illustrating all metabolites in the tricarboxylic acid cycle (TCA). Blue arrows indicate a decrease of these metabolites (highlighted in red) in *Tak1 CKO* mouse brains. Metabolites without arrows (black) exhibited no significant difference in concentration between Ctrl and *Tak1 CKO*. Relative abundance (normalized by TIC of the control group, y-axis) of the metabolites shown in E from control (light blue) and *Tak1 CKO* (magenta) samples. *, *p* < 0.05, **, *p* < 0.01, ***, *p* < 0.005; two-tailed unpaired Student’s *t*-test. FAD, flavin adenine dinucleotide. (**G**) Volcano plot representing significantly up- and down-regulated genes based on the log2(FC) and −log10(adjusted *p* value, *Padj*). The thresholds are FC>2 and *Padj*(FDR)<0.05 for Ctrl (n = 4 mice) and *Tak1 CKO* (n = 4 mice). Up-regulated and down-regulated genes are highlighted in pink and light blue, respectively; black vertical lines highlight FC of - 1.5 and 1.5, while black horizontal lines represents a *P*adj of 0.05. Core genes in the “Glutamatergic Synapse” geneset are in bold; blue represents genes with significant differential expression while brown represents genes with no significant difference. (**H**) GSEA plots of Glutamatergic Synapse geneset, with black bars indicating genesets represented among all genes pre-ranked by ranking metrics (Ctrl versus *Tak1 CKO*), with indicated normalized enrichment score (NES) and false discovery rate (FDR) q-value. (**I**) Color scale heatmap showing the normalized expression of core genes of the Glutamatergic Synapse geneset which is significantly down-regulated in the *Tak1 CKO* group vs Ctrl.

To further determine whether the abnormalities in mitochondria affected metabolic pathways involved in energy generation in the brain, we collected tissues from the cerebral cortex of control (WT or *Tak1^fl/fl^*) and *Tak1 CKO* mice for targeted metabolomic analyses and RNA sequencing. We measured over 200 metabolites for the metabolomic analysis, including 20 amino acids and their derivatives. These measurements covered most metabolites of the classical metabolic pathways in cells (Xiong et al., 2020). The metabolite profiles from the control and *Tak1 CKO* brains tended to cluster separately in unsupervised principal component analysis (PCA) (**Figure 7C**), indicating dramatic alterations in metabolomes between control and *Tak1 CKO* brains (**Figure 7C**). Among the metabolites that we detected, pyruvate, α-ketoglutarate (α-KG), fumarate, and NAD were significantly decreased in *Tak1 CKO* brains (**Figure 7D–F**). These metabolites are crucial for the TCA cycle and energy generation pathways, which is consistent with the morphological abnormality that we observed in mitochondria. In addition, we detected the glutamate concentration was much lower in *Tak1 CKO* compared to the control group (69.1±5.9%, n = 5 in *Tak1 CKO* brain. The concentration was normalized to that of the control group, **Figure 7D–F**). Our further RNA sequencing analysis showed that a gene cluster pertaining to glutamatergic transmission was downregulated in *Tak1 CKO* mice. The expression levels of the genes (e.g., *Plcb2, Gnas, Plcb3, Plas2g4a*, etc.; **Figure 7G** and **H**) associated with the glutamatergic synapse dramatically decreased (**Figure 7G–I**). Our results from both transcriptome and metabolomic measurements, as well as EM imaging, demonstrated that increasing vessel regression led to an alteration of the neuronal metabolism (i.e., energy generation and glutamate metabolism), and as a result decreased the activity in neurons. Given that it is extremely difficult for us to isolate the contribution of energy production and synaptic transmission to the reduction of neuronal activity in *Tak1 CKO* mice, further investigation with new methods/tools is required to continue this study in the future.

## Discussion

The term “brain plasticity” usually refers specifically to neuroplasticity because neurons are the most functionally relevant cell type in the brain (Kolb and Whishaw, 1998). Our data indicate that the blood-vessel pattern in the mature adult mammalian brain changes substantially with age. VEGF is a critical molecule for both vessel development and regression(Baffert et al., 2006). In the human brain, the responsiveness of HIF1 to hypoxia wanes with age, thereby reducing *VEGF* expression (Frenkel-Denkberg et al., 1999; Rivard et al., 2000). Our live-imaging experiments only rarely revealed new vessel formation in the adult mouse brain under physiological conditions. Accumulation of regressive vessels (our estimate: ~20% per year in mouse brain) might explain the dramatic reduction of blood vessels in the adult mammalian brain.

The distance over which oxygen can diffuse in the normal brain is highly regulated. The limit of oxygen diffusion is 100–150 μm in live tissue(Baish et al., 2011; Helmlinger et al., 1997). The distance between capillaries is ~40 μm in mice(Nicholson, 2001). In the white matter of the human brain, the capillary density is much lower than in the gray matter, and the inter-capillary distance can reach 100 μm. Vessel regression leads to a decrease in blood-vessel density and causes the inter-capillary distance to exceed the limit of oxygen diffusion, and the consequent reduction of oxygen availability to brain cells may result in stress on neurons or glial cells during periods of increased activity in a particular brain region; this may explain why the chronic low-grade ischemia caused by hyaline arteriolosclerosis (hypertension-induced disease of brain arterioles) or chronic cerebral hypoperfusion preferentially damages the white rather than the gray matter (Hatazawa et al., 1997), ultimately resulting in a diminution of the vasculature with consequent dementia.

It has been reported that alteration of neuronal activity causes remodeling of blood vessel patterns in the brain (Lacoste et al., 2014). Both endothelial cells and pericytes play an active role in the functional coupling of the neurovascular unit, in which blood vessels are the key component (Chen et al., 2014; Chow et al., 2020; Hall et al., 2014). Any dramatic alterations (remodeling, regression, angiogenesis, etc.) in the pattern of the brain vasculature may result in altered regulation of brain microcirculation, which in turn is likely to lead to changes in the local supply of glucose and oxygen in the brain (Abbott et al., 2006; Kaplan et al., 2020; Zlokovic, 2011). Neurons are the key player in brain function, and they consume the most glucose and oxygen in the adult mammalian brain (Belanger et al., 2011). After an alteration to the microcirculation, especially a decrease in blood supply followed by vessel regression, neurons will be the most vulnerable cell type under these stressed conditions, and they will sustain damage (Magistretti and Allaman, 2015). Although we detected a substantial decrease of neuronal activity in the cerebral cortex in *Tak1 CKO* mice due to an increasing accumulation of regressive vessels, further work is required to determine how blood vessel regression regulates energy-related metabolism under physiological conditions and whether a certain threshold of vessel regression leads to neuronal or synaptic degeneration.

## Supplemental Figure legends

**Figure S1.**
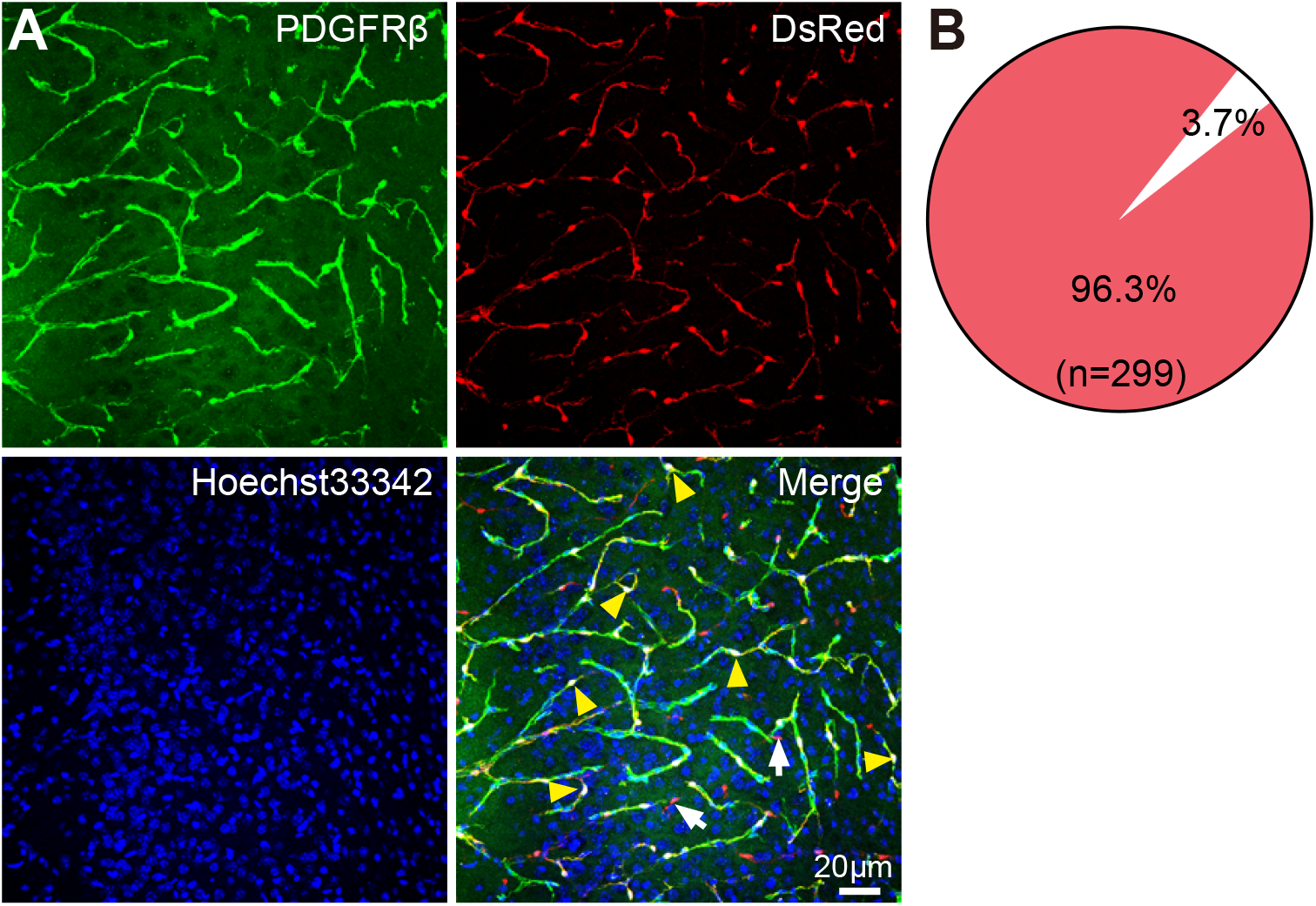
Nearly all pericytes in *NG2DsRedBACtg* mice are labeled with DsRed. (**A**) Brain sections from a *NG2DsRedBACtg* mouse labeled with anti-PDGFRβ. Red, DsRed fluorescence; blue, DNA nuclei, Hoechst 33342. (**B**) Nearly all (96.3%, n = 288 of 299 cells) pericytes (PDGFRβ^+^) expressed the DsRed protein. Arrowheads indicate cells that expressed DsRed and also were labeled with PDGFRβ. Arrows, NG2 glial cells expressing DsRed but not labeled with anti-PDGFRβ.

**Figure S2.**
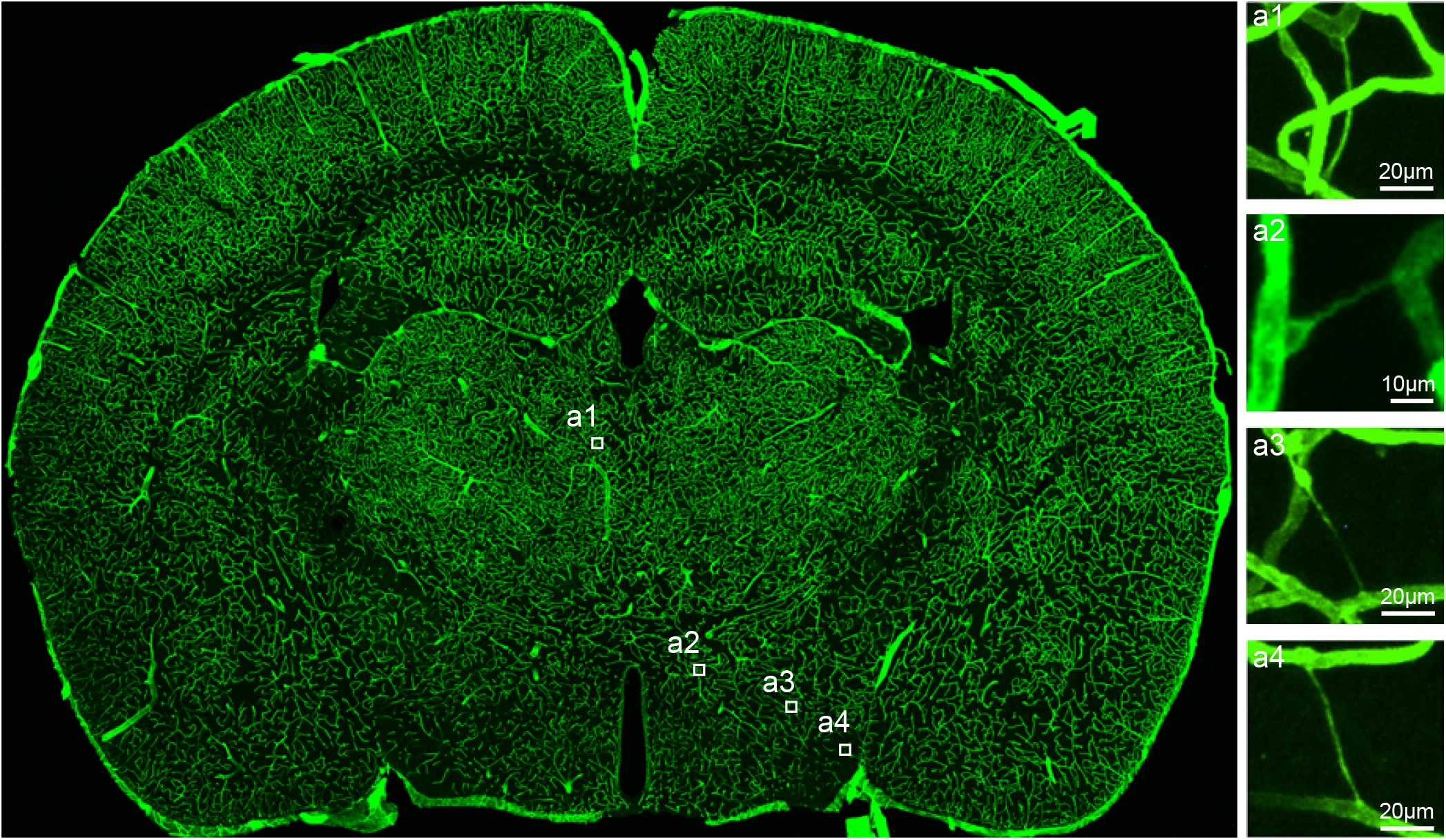
High-resolution images of all blood vessels from a whole section of an adult mouse brain. High-resolution image of a whole-brain section (60 *μ*m) from an adult mouse (P107). Insets 1–4 present four examples of regressing blood vessels in the brain section. The laminin layer of blood vessels was labeled by anti-collagen IV.

**Figure S3.**
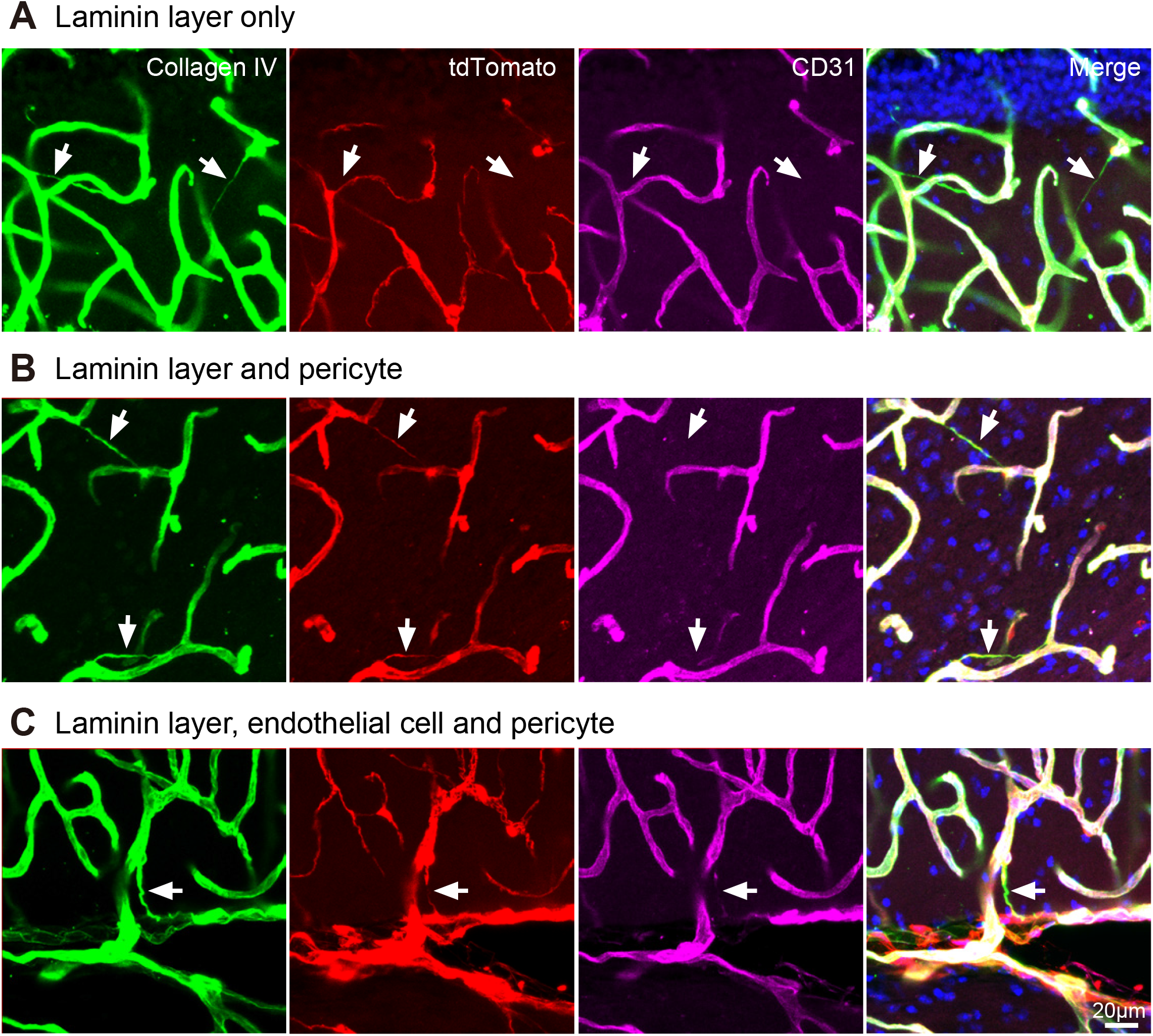
Characterization of the cellular components of regressive vessels in the brain of *Pdgfrβ-Cre::Ai14* mice. Three different regressing vessels were observed in a section of a *Pdgfrβ-Cre::Ai14* mouse brain after staining with anti-CD31 and anti-collagen IV. Some regressive vessels had laminin layers but were devoid of endothelial cells or pericytes (**A**). Other regressive vessels had both laminin layers and pericytes (**B**). Still other regressive vessels had endothelial cells, pericytes, and laminin layers. Each laminin layer was stained with anti-collagen IV (green, **C**), endothelial cells were labeled with anti-CD31 (purple), and pericytes were labeled with tdTomato (red), and the nuclei were labeled with Hoechst 33342 (blue).

**Figure S4.**
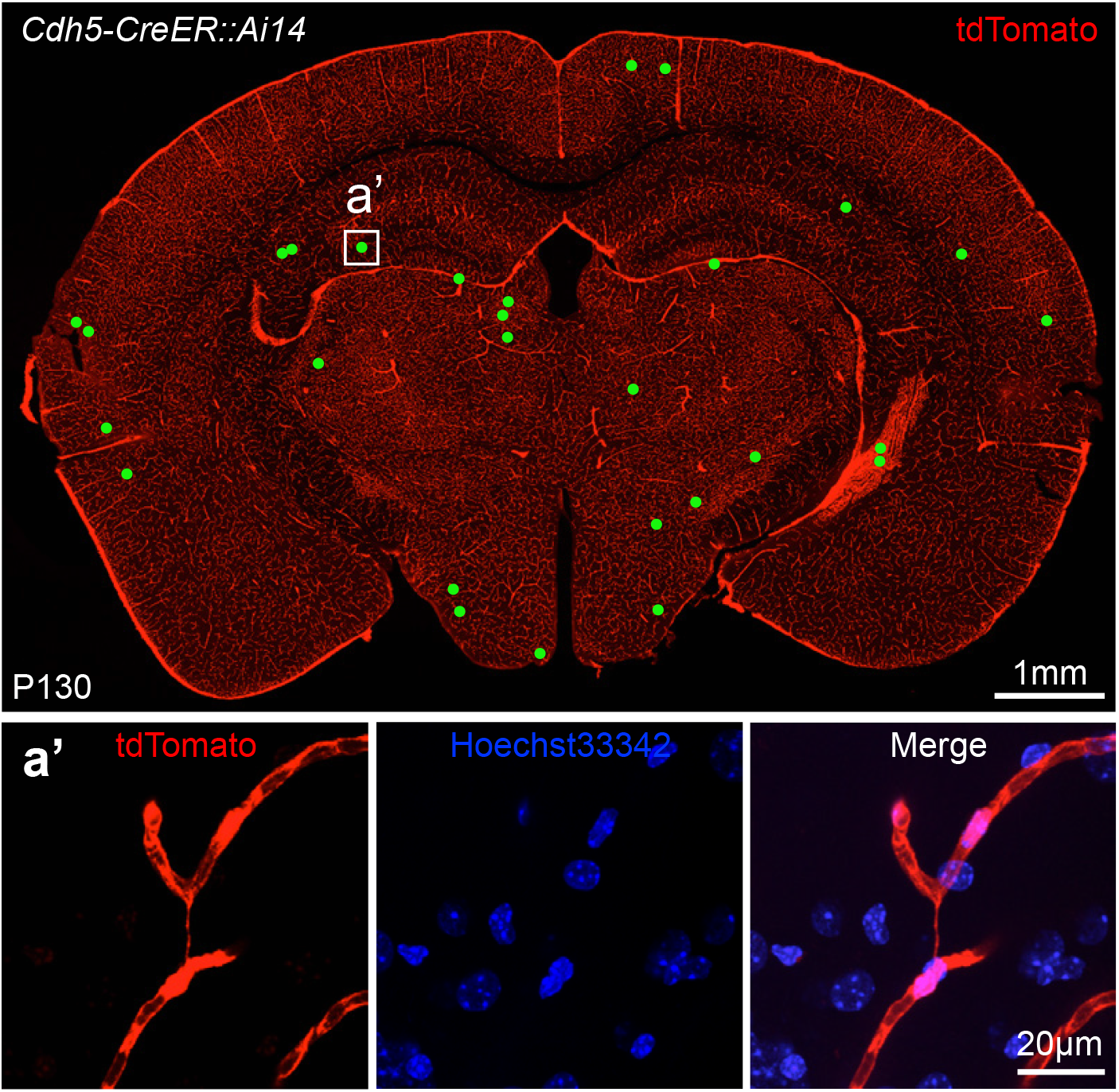
Distribution of regressing vessels containing endothelial cell components in *Cdh5-CreER::Ai14* mice. High-resolution image of a whole-brain section from a *Cdh5-CreER::Ai14* mouse (P78). Green dots indicate each regressive vessel with endothelial cell components. a’, Inset presents a representative regressive vessel containing components of endothelial cells. Red, tdTomato signal from endothelial cells; blue, nuclei stained with Hoechst 33342.

**Figure S5.**
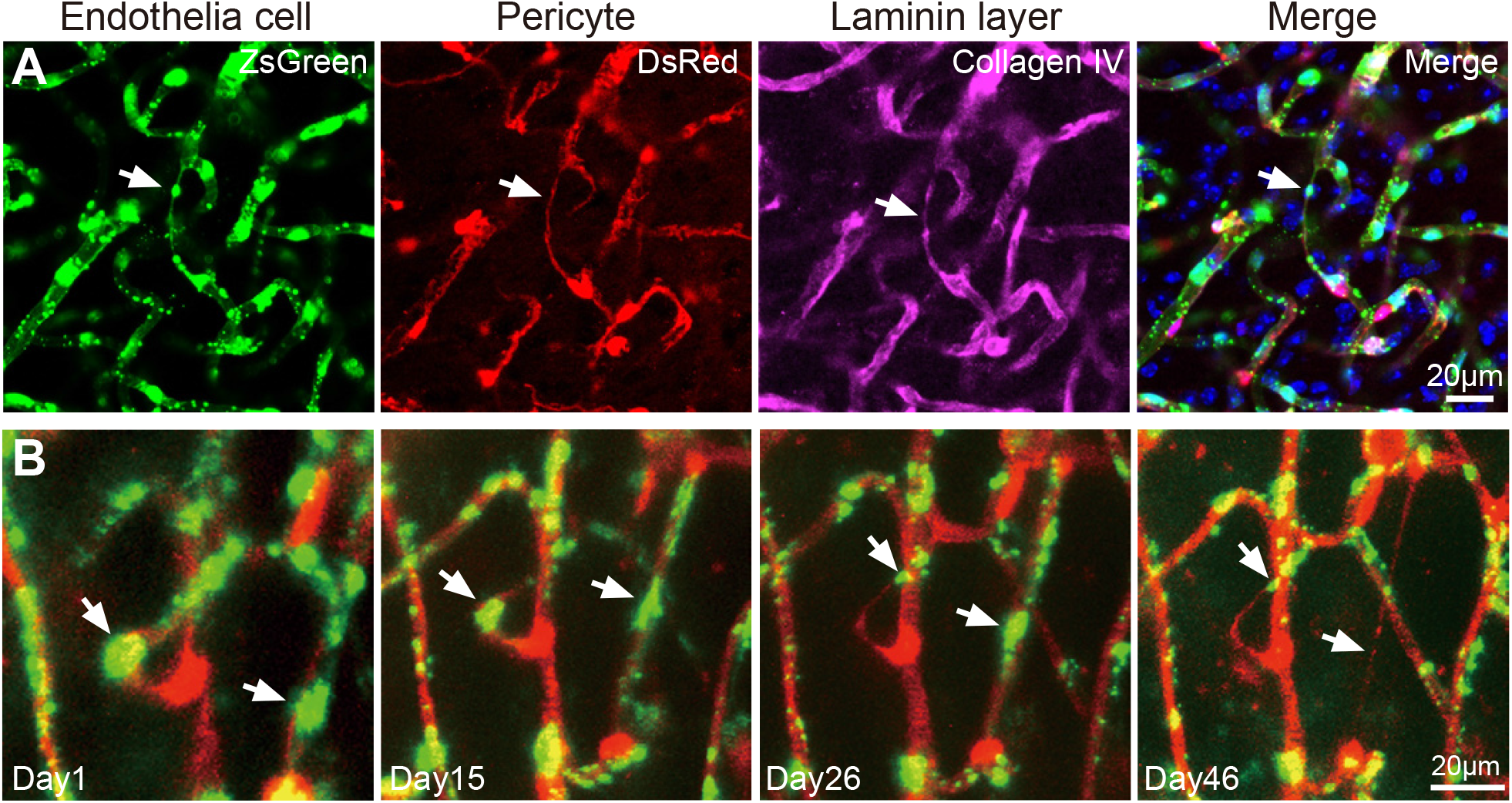
Time-lapse *in vivo* imaging of a regressing vessel the brain of a *Cdh5-CreER::Ai6::NG2DsRedBAC* mouse. (**A**) Regressive blood vessel stained with anti-collagen IV in a *Cdh5-CreER::Ai6::NG2DsRedBAC* mouse. Purple, signal from Alexa 647 after staining the laminin layer with anti-collagen IV. Arrows, one regressive vessel. (**B**) Endothelial cells (green, ZsGreen from *Cdh5-Cre::Ai6*), pericytes (red, DsRed from *NG2DsRedBAC*), and blood flow (also red signal) were simultaneously imaged. Arrows indicate two regressing vessels at different time points (days 1, 15, 26, and 46).

**Figure S6.**
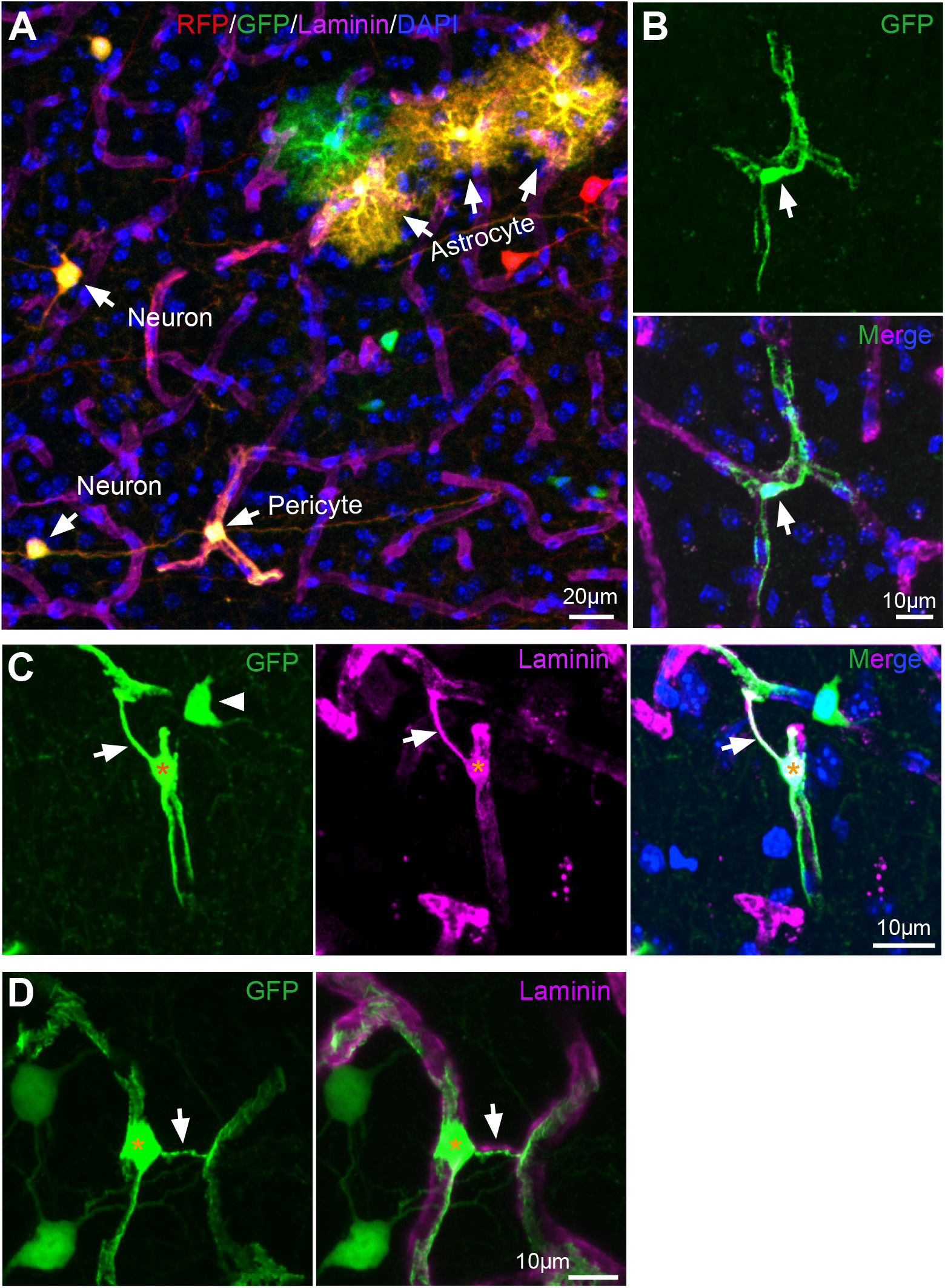
Morphology of pericytes in regressing vessels. (**A**) An example image of a brain section of a *Hprt-Cre::MADM* mouse. Neurons and glial cells are labeled, and pericytes were sparsely labeled (arrows). (**B**) Morphology of a typical pericyte. (**C, D**) Gross morphology of an individual pericyte from a regressing vessel in brain of a *Hprt-Cre::MADM* mouse. Arrows, regressing vessels. Asterisks indicate the somas of the pericyte in regressing vessels. The processes extending from pericytes on both blood vessels were very complex, indicating a stable structure. A neuron (arrowhead) is close to the pericyte in (C). Purple, signal from anti-laminin; blue, nuclei labeled with Hoechst 33342; red, red fluorescence protein (RFP); green, GFP.

## Legends for Videos

**Video 1**

**Time-lapse *in vivo* imaging of blood microcirculation.**

Time-lapse *in vivo* imaging of blood microcirculation in the cerebral cortex of an adult *NG2DsRedBAC* transgenic mouse 7 days before occlusion. Green, FITC-dextran 500K; red, DsRed fluorescence from pericytes of blood vessels.

**Video 2**

**Time-lapse *in vivo* imaging of blood microcirculation.**

Time-lapse *in vivo* imaging of blood microcirculation in the cerebral cortex of an adult *NG2DsRedBAC* transgenic mouse 1 day after occlusion. Green, FITC-dextran 500K; red, DsRed fluorescence from pericytes of blood vessels. One blood vessel (arrow) was already occluded.

**Video 3**

**Time-lapse *in vivo* imaging of blood microcirculation.**

Time-lapse *in vivo* imaging of blood microcirculation in the cerebral cortex of an adult *NG2DsRedBAC* transgenic mouse 11 days after occlusion. Green, FITC-dextran 500K; red, DsRed fluorescence from pericytes of blood vessels. The occluded blood in Extended Data Video 2 (1 day after occlusion) had already regressed (Video 3). Very dim DsRed and FITC fluorescence detected in the regressive vessel.

**Video 4**

**Time-lapse *in vivo* imaging of blood microcirculation**

Time-lapse *in vivo* imaging of blood microcirculation in the cerebral cortex of an adult *NG2DsRedBAC* transgenic mouse 27 days after occlusion. Green, FITC-dextran 500K; red, DsRed fluorescence from pericytes of blood vessels. No DsRed and FITC fluorescence were detected in the regressive vessel that was occluded in Extended Data Video 2.

**Video 5**

**The distribution of all regressive vessels in one brain section.**

Images showing the distribution of all regressive vessels taken from a 500-*μ*m brain section after it was stained with anti-collagen IV and then cleared using the PEGASOS method. Green, fluorescence from anti-collagen IV. All of the regressive vessels in this optical block (1380 *μ*m × 1380 *μ*m × 248 *μ*m) are highlighted with red pseudo-color added using Imaris9.0 software.

**Video 6**

**Calcium activity in a mouse brain of *Cdh5-CreER:Tak1^fl/fl^*.**

Example video of the GCaMP6 signal obtained from a *Cdh5-CreER:Tak1^fl/fl^* (Tak1 CKO) mouse brain before administration of tamoxifen (frequency 5 Hz, for 10 min).

**Video 7**

**Calcium activity in a mouse brain of *Cdh5-CreER:Tak1^fl/fl^*.**

Example video of the GCaMP6 signal obtained from a *Cdh5-CreER:Tak1fl/fl* (Tak1 CKO) mouse brain 1 week after administration of tamoxifen (frequency 5 Hz, for 10 min).

**Video 8**

**Calcium activity in a mouse brain of *Tak1^fl/fl^*.**

Example video of the GCaMP6 signal obtained from a *Tak1fl/fl* (Tak1 Control) mouse brain before administration of tamoxifen (frequency 5 Hz, for 10 min).

**Video 9**

**Calcium activity in a mouse brain of *Tak1^fl/fl^*.**

Example video of the GCaMP6 signal obtained from a *Tak1^fl/fl^* (Tak1 Control) mouse brain after administration of tamoxifen (frequency 5 Hz, for 10 min).

## ACKNOWLEDGEMENTS

The project was initiated in the Jan lab at UCSF. We thank Lily Jan and Yuh-Nung Jan’s generous support. We thank Liqun Luo’s lab for providing MADM-7 mice and Rolf A Brekken for VEGF-antibodies. Drs. Yuanquan Song (UPenn), Zhaozhu Hu (JHU), Ji Hu (ShanghaiTech), Yang Xiang (U. Mass), Hao Wang (Zhejiang U.) and Ruikang Wang (U. Washington) for critical input, colleagues at Children’s Research Institute, Departments of Neuroscience, Neurology and Neurotherapeutics, Pediatrics from UT Southwestern, and colleagues from the Jan lab for discussion. Dr. Bridget Samuels, Sean Morrison (UT Southwestern), and Nannan Lu (Zhejiang U.) for critical reading. We acknowledge the assistance of the CIBR Imaging core. We also thank UT Southwestern Live Cell Imaging Facility, a Shared Resource of the Harold C. Simmons Cancer Center, supported in part by an NCI Cancer Center Support Grant, P30 CA142543K. This work is supported by CIBR funds and the American Heart Association AWRP Summer 2016 Innovative Research Grant (17IRG33410377) to W-P.G.; National Natural Science Foundation of China (No. 81370031) to Z.Z.; National Key Research and Development Program of China (2016YFE0125400) to F.H.; National Natural Science Foundations of China (No. 81473202) to Y.L.; National Natural Science Foundation of China (No.31600839) and Shenzhen Science and Technology Research Program (JCYJ20170818163320865) to B.P.; National Natural Science Foundation of China (No. 31800864) and Westlake University start-up funds to J-M. J. NIH R01NS088627 to W.L.J.; NIH: R01 AG020670 and RF1AG054111 to H.Z.; R01 NS088555 to A.M.S., and European Research Council No.725780 to S.H.; W-P.G. was a recipient of Bugher-American Heart Association Dan Adams Thinking Outside the Box Award.

## AUTHOR CONTRIBUTIONS

W-P.G. and X.G. conceived the project. X.G. and W-P.G. designed experiments. X.G. performed all *in vivo* imaging experiments in animals and analyzed data. X.G., W-P.G., B.C., X-J C, J-L.L., N.L., Y. Y., F.C., J-M.J., B.S., Y.Y., S.Z., Y-C. S., K.S., N.E.P., Y.H., K.P., Z.Z., D.B.B., S-b.Y., W.S. performed the other experiments. W-P.G. R.H.A., V.L., L-J.W, B.P., H.Z., B.O.Z., F.H., S.H., A.M.S., M.M., X.W., Q.L. Y-m L, J. K., M. T., H. L., M. P., R.M.B. and H.Z. provided reagents and discussed experimental plans and the results. W-P.G. and X.G. wrote the manuscript. All authors reviewed and edited the manuscript.

## METHODS

### Animals

All animal experiments were conducted in accordance with protocols approved by the Institutional Animal Care and Use Committee at the University of Texas Southwestern Medical Center. MADM mouse strain was originally from Dr. Liqun Luo lab at Stanford (also available from Jackson lab, Cat# 021457). The GT/TG locus is located on Ch7. *Hprt-Cre* mouse line was from Jackson lab (Cat# 004302). *Tak1^fl/fl^* mouse line was purchased from Jackson lab (Cat# 011038). *NG2DsRedBAC* was originally from Akiko Nishiyama lab (also available at Jackson lab, Cat# 008241). *Pdgfrb-Cre* from Volkhard Lindner’s lab. *Cdh5-CreER* was from Ralf H. Adams’s lab. *Ai14* and *Ai6* is available from the Jackson lab (Cat# 007908 for *Ai14* and 007906 for *Ai6*).

### *In vivo* labeling of brain microcirculation

Fluorescein isothiocyanate-Dextran 500,000-Conjugate (FITC-Dextran, Sigma Aldrich) or TRITC-dextran (Sigma Aldrich) was prepared in saline (0.9% NaCl) at a concentration of 10mg/ml. Adult mice were anesthetized with a mixture of ketamine (80–100 mg/kg) and xylazine (10–12 mg/kg). The tail was warmed with a heat lamp for about 1min and then wiped with 70% ethanol around the injection site. A 31G insulin syringe needle was inserted with the bevel up, at an angle of 5–15 degrees into the vein. 80–100*μ*l FITC-Dextran 500K solution was injected. Blood circulation could then be detected via the FITC signal, and blood cells were visualized by contrast as dark areas without fluorescent signal.

### Fluorescent labelling of blood cells

Labelling was conducted as previously reported with some modifications(Jia et al., 2017). Briefly, 30–100μl of blood (2–4 drops) was taken from the submandibular veins of the mouse after poking its cheek with a Goldenrod animal bleeding lancet. The blood was collected in an Eppendorf tube containing 500 μl of 1× Hank’s Balanced Salt Solution (HBSS) with 10 mM EDTA added as an anticoagulant. The whole blood was centrifuged at 150 g for 3 min. After the supernatant was removed, the pellet was resuspended in 1ml HBSS containing DiO (7 μM, Thermofisher Scientific). Cells were then incubated at 37°C for 15 min. Labelled blood cells were washed twice with HBSS and then centrifuged and resuspended in 300 μl HBSS. About 50μl solution with DiO-labeled cells was injected back to the same mouse through the tail vein. DiO dye was excited with a laser at 488nm or 860nm IR-laser on Zeiss LSM710.

### Longitudinal time-lapse imaging of brain vasculature *in vivo*

Glass cranial windows were made in the skulls of different transgenic mice, allowing 1–2 months recovery from the craniotomy surgery before we performed live imaging on an upright Zeiss LSM710 NLO two-photon excitation microscope with a 20×/1.0 water-immersion objective lens (Zeiss). Regions of interest (ROI) were scanned with XYZ mode for time-lapse imaging as we previously reported(Ge et al., 2012; Jia et al., 2017). The same ROI area was re-located using the branching pattern of major blood vessels, and Z-stack images were scanned once a week (1–2 h) in the following 1–6 months. We measured the speed of blood flow in the visualized vessels with the line scanning function. During imaging, mice were anaesthetized with 1–2% isoflurane in oxygen, and their body temperature was kept with a home-made heating pad (10 × 5 cm). Blood vessels were visualized by FITC–dextran injected through mouse the tail vein in *NG2DsRedBAC* tg mice. Blood vessels in the brains of *Cdh5-CreER::Ai6::NG2DsRedBAC* triple transgenic mice were visualized by TRITC-dextran. DsRed, FITC, or ZsGreen was excited with IR laser (930-960 nm for DsRed, 860nm for FITC/ZsGreen) or one photon laser (543 nm for DsRed, 488nm for FITC/ZsGreen).

### *In vivo* imaging of neuronal activity in conscious mice

AAV-Camk2-GCaMP6m (in 0.5 μl phosphate-buffered saline with a titer of 2.4 × 10^12^) was injected into layers II–IV of the cerebral cortex of mice at the age of 8–10 weeks. A glass cranial window was built using the same surgery procedure described above for longitudinal time-lapse imaging. We usually measured the GCaMP6 signal approximately 20–30 days post-injection of AAV from layers II–IV around the injection site in *Cdh5-CreER::Tak1^fl/fl^* mice. Tamoxifen (Sigma, Cat# T5648) was administered intraperitoneally (70 μg/kg body weight; 10 mg/ml tamoxifen was dissolved in mixed solution of corn oil and ethanol, 9:1 v/v). A two-photon laser (950 nm) was used to image GCaMP transients 200–700 μm below the pia in conscious mice with a 20× water immersion lens (N.A., 1.0) mounted in a Zeiss LSM780 microscope (setting, 512 × 512, frequency 5 Hz, for 10 min). Images were processed with Image J to generate the ΔF/F curve as described(Barson et al., 2020). Briefly, the fluorescence density of each ROI (region of interest) time series was measured, with the baseline fluorescence (*F_0_*) being defined as the average of the lowest 10% of samples. The instantaneous fluorescence of the ROI time series is F, and ΔF = F – F_0_. The 100 × ΔF/F curve was plotted using clampfit 10.0 software. The frequency of GCaMP-mediated calcium transients was detected with MiniAnalysis software. Quantitative data were processed with GraphPad Prism software, and statistical analysis was carried out with the Student’s *t*-test.

### Immunohistochemistry

Mice of different ages were perfused transcardially with 20 ml of PBS, followed by 20 ml of 4% (w/v) paraformaldehyde (PFA) in PBS. The brain was fixed in cold PFA in 4°C for 1.5–2 h, washed with a large volume of PBS overnight in 4°C, and dehydrated with 15% and 30% sucrose in PBS sequentially. The human brain was fixed whole in 20% formalin for 9 days and then dissected, showing no gross abnormalities; samples were taken from the frontal lobe cortex with subcortical white matter, cerebellum, hippocampus, and basal ganglia (caudate and putamen). The decedent was a 45-year-old man who died from the complications of disseminated neurodendocrine carcinoma of the pancreas; the permission to use tissue for research was covered by the autopsy permit signed by his next-of-kin. Fixed mouse brains and monkey or human brain samples were sectioned with a cryostat (model CM3050S, Leica) or a vibratome (Leica) into sections of 20-70 μm thickness. Sections were permeabilized with 0.25% Triton X-100 and then blocked with 5% BSA and 3% normal goat serum with 0.125% Triton X-100 for 2 h. Primary antibodies against mouse or human laminin (1:300 rabbit, Sigma), mouse collagen IV (EMD Millipore), mouse CD31 (1:300, rat, Abcam), or mouse PDGFRβ/CD140b (1:300, rat, eBioscience) were incubated with brain sections for 24-48 h at 4°C. Together with Hoechst33342 or DAPI (1 μg/ml), secondary antibodies conjugated with Alexa488, 546 or 647 (1:500, Life Technologies) were used after 2 h incubation at RT (22-25°C). Sections were mounted with anti-fade mounting medium Fluoro-Gel (EMS). All images were taken with a Zeiss LSM710 NLO confocal microscope. Whole-section images were scanned with the tiling function on an inverted Zeiss LSM780 confocal microscope, with a 20×/0.8 air objective lens or 63×/1.4 oil objective lens.

### Short-term occlusion of middle cerebral artery (MCA)

Mice (>P100) were anesthetized with isoflurane until unconscious followed by 1.5-2% isoflurane during MCA occlusion surgery as described in our previous study (Jia et al., 2017) and body temperature was maintained during surgery with a homemade heating pad (10 × 5 cm). A midline neck incision was made, the left common carotid artery (CCA) was carefully separated from the vagus nerve, and the artery was ligated using a 5.0-string. A second knot was made on the left external carotid artery (ECA). The left internal carotid artery was isolated, and a knot of a 5.0 string was left loose. This knot was not tightened until the intraluminal insertion was done. A small hole was cut in the CCA before it was bifurcated to the ECA and ICA. A silicon-coated monofilament (tip diameter = 230 *μ*m for adults, Doccol Corporation) was inserted into the ICA until it stopped at the origin of the MCA in the circle of Willis. The third knot on the ICA was closed to fix the filament in position. During MCA occlusion (15 and 20 min), mice were kept under anesthesia on the heating pad. After occlusion, the knot on the ICA was loosened and the filament was then removed. The knot on the ICA was later tightened again, and the midline neck incision was sewn closed. The mice were put back into their home cages to allow the blood reperfusion in the occluded brain hemisphere for 1 or 2 weeks.

### Tissue clearing with PEGASOS

Thick slices were cleared as previously (Jing et al., 2018). Briefly, immunostaining (1^st^ antibody, anti-collagen IV, 1:300, incubation time, 7d; 2^nd^ antibody, Alexa 488, 1:500, incubation time, 3d at 4°C) was performed before we started the tissue clearing. Brain slices (500μm) were then treated with Quadrol decolorization solution for 2 d at 37 °C after fixation with 4% PFA solution for 24 h. Samples were then immersed in gradient delipidation solutions in a 37 °C shaker for 1 to 2 d, followed by dehydration solution treatment for 1 to 2 d and BB-PEG clearing medium treatment for at least 1 d until reaching transparency. Slices were then preserved in the clearing medium at RT before imaging with a confocal microscope (Zeiss LSM780).

### Image data analysis

Image data analysis was done with Image J software (NIH) and ZEN image processing software from Zeiss. The number of regressing vessels (RV) was counted manually with ZEN software. RV density per square millimetre was normalized from the optical volume of the images. The blood vessel length density (Figure 3) was measured by calculating the area of all blood vessels from 3-D projection images in Image J. 3D reconstruction and movies of 3D blood vessels and RV distribution (Extended Data Video 5) were produced with Imaris 9.0 (Bitplane).

### Ly6G antibody treatment in LPS-treated mouse

Ly6G antibody (BD Pharmingen, #551459) was administered intraperitoneally (i.p.) at a dosage of 4mg/kg b.w. for the 1st injection and 2mg/kg b.w. for the following injections. The control antibody was also administered i.p. with the same concentration. Treatments were performed on 12–14-week-old C57BL/6J mice on days 1, 4, 7, 10, 13, 16, and 19 of the study. The injection of the antibodies was combined with the 1-week LPS injection (0.5μg/g b.w., 4 times) schedule as described above (Figure 5A). Mouse brains were collected on day 21 for staining.

### Imaging mitochondria with electron microscopy

The subcellular mitochondrial morphology and synaptic structure were imaged using electron microscopy (EM). On days 7-10 after tamoxifen injection at 12-14 weeks old, *Cdh5-CreER::Tak1^fl/fl^* and their littermates (control mice) were anesthetized and transcardially perfused with EM fixation solution (4% PFA in 0.1M sodium cacodylate buffer, pH7.4, containing 0.1% glutaraldehyde) at room temperature. The brain was collected and then fixed in solution (2.5% glutaraldehyde in 0.1M Sodium Cacodylate Buffer, pH7.4) for 2h at 4°C. 1 mm cubic blocks of cortical tissue and hippocampus tissue were collected and embedded in resin embedding medium (Epon). Blocks were sectioned with a diamond blade (Diatome) on a Leica Ultracut 7 ultramicrotome (Leica Microsystems) and collected onto copper grids. Thin sections were negatively stained with 2% aqueous uranyl acetate. Images were acquired using a Morada Digital Camera and iTEM software (Olympus) under an FEI Tecnai G2 Spirit Biotwin transmission electron microscope, at voltage of 120 KV. Quantitative data and statistical analysis were processed in Graphpad Prism 7.01 software and ImageJ.

### Purification of metabolites from brain tissue

Cortical tissue samples from the rostral hemisphere (1/2 of the cortical region of the whole hemisphere) were collected in an Eppendorf tube and then 2ml ice-cold methanol/80% water (vol/vol) was added before weighing. The samples were homogenized and vortexed for 5min. We transferred the extracted solution with 5mg brain tissue to 900*μ*l 80% ice-cold methanol/80% water (vol/vol) and vortexed for 1 min. After centrifugation at 17,000 g for 15 min at 4°C, 800 μl of the supernatant was transferred to a new tube. All samples were evaporated until dry using a SpeedVac concentrator (Thermo Savant). The samples were stored under −80 °C before we performed targeted metabolomic measurement.

### Targeted Metabolomics and data analysis

Metabolites extracted from the cerebral cortex were reconstituted in 50 *μ*l 0.03% formic acid in water and then analyzed with a SCIEX QTRAP 5500 liquid chromatograph/triple quadrupole mass spectrometer as done in our previous study(Xiong et al., 2020; Yu et al., 2020). Using a Nexera Ultra-High-Performance Liquid Chromatograph system (Shimadzu Corporation), we achieved separation on a Phenomenex Synergi Polar-RP HPLC column (150 × 2 mm, 4 *μ*m, 80 Å). The mass spectrometer was used with an electrospray ionization (ESI) source in multiple reaction monitoring (MRM) mode. We set the flow rate to 0.5 ml/min and the injection volume 20 *μ*l. We acquired MRM data with Analyst 1.6.3 software (SCIEX).

### Metabolomics data analysis

Integrated chromatogram peaks of each metabolite were analyzed with MultiQuant software (AB Sciex). The ion intensity was calculated by normalizing single ion values against the total ion value of the entire chromatogram (i.e. Total Ion Chromatogram/TIC)(Yu et al., 2020). The data matrix was input into SIMCA-P software (Umetrics) by mean-centering and Pareto scaling for subsequent analysis so that the model fitting would not be biased by concentrations and variations of different metabolites(Xiong et al., 2020). Both unsupervised and supervised multivariate data analysis approaches including principal component analysis (PCA) were performed using Metaboanalyst 4.0 (Chong et al., 2018). All data are presented as mean ± s.e.m.

### RNA sequencing (RNA-Seq) and differential expression analysis

*Cdh5creER::Tak1^fl/fl^* mice and their littermate control mice at the age of 12–14 weeks were treated with one dose of tamoxifen (700mg/kg b.w.) via intraperitoneal injection (i.p.). The cerebral cortex was freshly collected on dpi7 for total RNA extraction (RNeasy Mini kit, Qiagen #74104). RNAseq library was prepared by DNA SMART ChIP Seq Kit (TAKARA #101617). RNA Sequencing was performed on Illumina NextSeq 500 desktop Next Generation Sequencing (NGS) system. Sequencing reads were aligned to mouse reference genome GENCODE Version M9. Differentially expressed genes were identified by normalized ratio of Reads Per Kilobase of transcript per million mapped reads (RPKM) between the *Tak1 CKO* and littermate control mice.

Fastp (Chen et al., 2018) was recruited to low-quality reads and adaptor trimming with a default setting (Bolger et al., 2014) (http://www.usadellab.org/cms/?page=trimmomatic). Cleaned reads were mapped to the ensembl mouse reference genome GRCm38.p6 (http://asia.ensembl.org/Mus_musculus/Info/Index) using STAR alignment software (Dobin et al., 2013). The mapped reads were counted to genes using featureCounts (http://subread.sourceforge.net/). Differential expression analysis was performed using DESeq2 (https://bioconductor.org/packages/release/bioc/html/DESeq2.html) with a cutoff of FDR < 0.05 and abs (log2FC)>1. Volcano plot, Scatter plotting and heatmaps were generated using R packages (ggplot2; pheatmap) implemented in R studio.

### GO and KEGG enrichment analysis

Functional enrichment in GO terms (Cellular Component; Biological Process; Molecular Function) of differential expression genes (FDR<0.05 & |FC| >2) was performed using the clusterProfiler R package (Yu et al., 2012), setting a q value threshold of 0.05 for statistical significance.

### GSEA analysis

The mechanisms underlying the relationship between blood vascular regression and neuron activity depression were explored with GSEA (Reimand et al., 2019). For gene set enrichment analysis (GSEA), we generated a KEGG_2019_Mouse geneset based on a database file from Enrichr online library (https://amp.pharm.mssm.edu/Enrichr/)(Chen et al., 2013). Genes were pre-ranked through the metrics algorithm (we applied sign of log fold change * −log10(p-value[not adjusted p-val)]; statistical result of DESeq2. Pre-ranked (.rnk) file and custom geneset were used as input for GSEA v4.0.3 (https://www.gsea-msigdb.org/gsea/index.jsp). The number of permutations was set at 1000 and enrichment statistics were set at “weighted”. For the general significance threshold, false discovery rate (FDR) q-val < 0.25 and |NES|>1.5 were considered as significant enrichment.

